# Mapping cell-cell fusion at single-cell resolution

**DOI:** 10.1101/2024.12.11.627873

**Authors:** Andrea L. Gardner, Lan Zheng, Kennedy Howland, Andrew Saunders, Andrea Ramirez, Patrik Parker, Chisom Iloegbunam, Daylin Morgan, Tyler A. Jost, Amy Brock

## Abstract

Cell-cell fusion is a tightly controlled process in the human body known to be involved in fertilization, placental development, muscle growth, bone remodeling, and viral response. Fusion between cancer cells results first in a whole-genome doubled state, which may be followed by the generation of aneuploidies; these genomic alterations are known drivers of tumor evolution. The role of cell-cell fusion in cancer progression and treatment response has been understudied due to limited experimental systems for tracking and analyzing individual fusion events. To meet this need, we developed a molecular toolkit to map the origins and outcomes of individual cell fusion events within a tumor cell population. This platform, ClonMapper Duo (‘CMDuo’), identifies cells that have undergone cell-cell fusion through a combination of reporter expression and engineered fluorescence-associated index sequences paired to randomly generated nucleotide barcodes. scRNA-seq of the indexed barcodes enables the mapping of each set of parental cells and fusion progeny throughout the cell population. In triple-negative breast cancer cells CMDuo uncovered subclonal transcriptomic hybridization and unveiled distinct cell-states which arise in direct consequence of homotypic cell-cell fusion. CMDuo is a platform that enables mapping of cell-cell fusion events in high-throughput single cell data and enables the study of cell fusion in disease progression and therapeutic response.

## Introduction

Intratumoral heterogeneity (ITH) is a defining feature of many tumor types which negatively affects prognosis^1–3^. Subclonal cancer cell populations within tumors may have unique properties which alter tumor aggressiveness and contribute to treatment failure. Consequently, ITH is especially problematic in cancers such as triple-negative breast cancer (TNBC) which do not have targeted treatment options and for which broad-spectrum chemotherapies remain the standard of care^4–6^. To develop more effective treatments for TNBC there is an urgent need to better understand the molecular drivers of ITH and tumor evolution.

Cell-cell fusion is a form of whole genome doubling (WGD), a known driver of ITH^7–13^. Novel methods have been developed to reconstruct WGD-aware subclonal phylogenies from patient data and more than 45% of breast cancers analyzed in the TCGA dataset show evidence of prior WGD^7–12^. While there is robust data to support WGD in tumor evolution and ITH generation, these phylogenetic reconstruction methods assume asexuality of tumor cells which may not be fully accurate^13,14^. Though understudied, the ability of cancer cells to fuse with other cells in the tumor microenvironment has been recognized for over a century^13,15–24^, with recent studies suggesting roles for cell-cell fusion in cancer stemness, metastatic potential, karyotype changes, and ITH^13,17,18,21,25,26^. Recent advancements in live-cell imaging have generated increased interest in cell-cell fusion, but no tools exist to study and track cell-cell fusion events in high-throughput.

To meet this need, ClonMapper Duo (“CMDuo”) was developed to allow clonally-resolved tracking of cell-cell fusion at single-cell resolution. In this study, we constructed high-resolution maps of cell-cell fusion events within two TNBC cell lines, identified unique cell states driven by cell-cell fusion, and linked transcriptomic features of parents to progeny at single-cell resolution.

## Results

### ClonMapper Duo is a two-color, two-index expressed DNA barcoding platform for tracking cell-cell fusion at single-cell resolution

CMDuo barcoded cells contain DNA barcodes that are transcribed as both polyadenylated transcripts and as CRISPR-compatible sgRNAs (Fig. 1a), allowing assignment to individual cells in scRNA-seq. To track cell-cell fusion using common live-cell imaging systems, CMDuo was engineered to co-express barcodes with nuclear-localized histone H2B-GFP (“CMDuo-green”) or H2B-mCherry (“CMDuo-red”) (Fig. 1a). Adding nuclear localization to the fluorescent signal allows precise cell counting and quantification from live cell imaging, and specific localization to H2B keeps fluorescence associated with parental nuclei in a shared cytoplasmic space until nuclear fusion takes place^27^ (Fig. 1b-c). The identification of fusion cells in barcoded cell populations is a challenge as cell doublets, common in many scRNA-seq preparations, or cells with multiple viral integrations also appear as multi-barcoded cells in scRNA-seq^28,29^. Given the rarity of fusion events compared to the rate of doublets in scRNA-seq^13,28,30^, potential fusion cells are likely filtered out and lost from analysis. To overcome this challenge, CMDuo was designed to confidently assign individual cells as fusion progeny in scRNA-seq using a combination of FACS enrichment on fluorescent identity and engineered fluorescence-associated index sequences paired to each random barcode sequence. For each fluorescent protein, four unique 5-nt barcode index sequences were designed by maximizing Hamming distance between all pairs^31^ (Supp. Fig. 1, Supp. Table 1). With the complete CMDuo platform, parental cells will express either GFP with a single CMDuo-green barcode or mCherry with a CMDuo-red barcode, while fusion cells will express both GFP and mCherry and contain both a CMDuo-green and CMDuo-red barcode (Fig. 1b-c).

**Figure 1.**
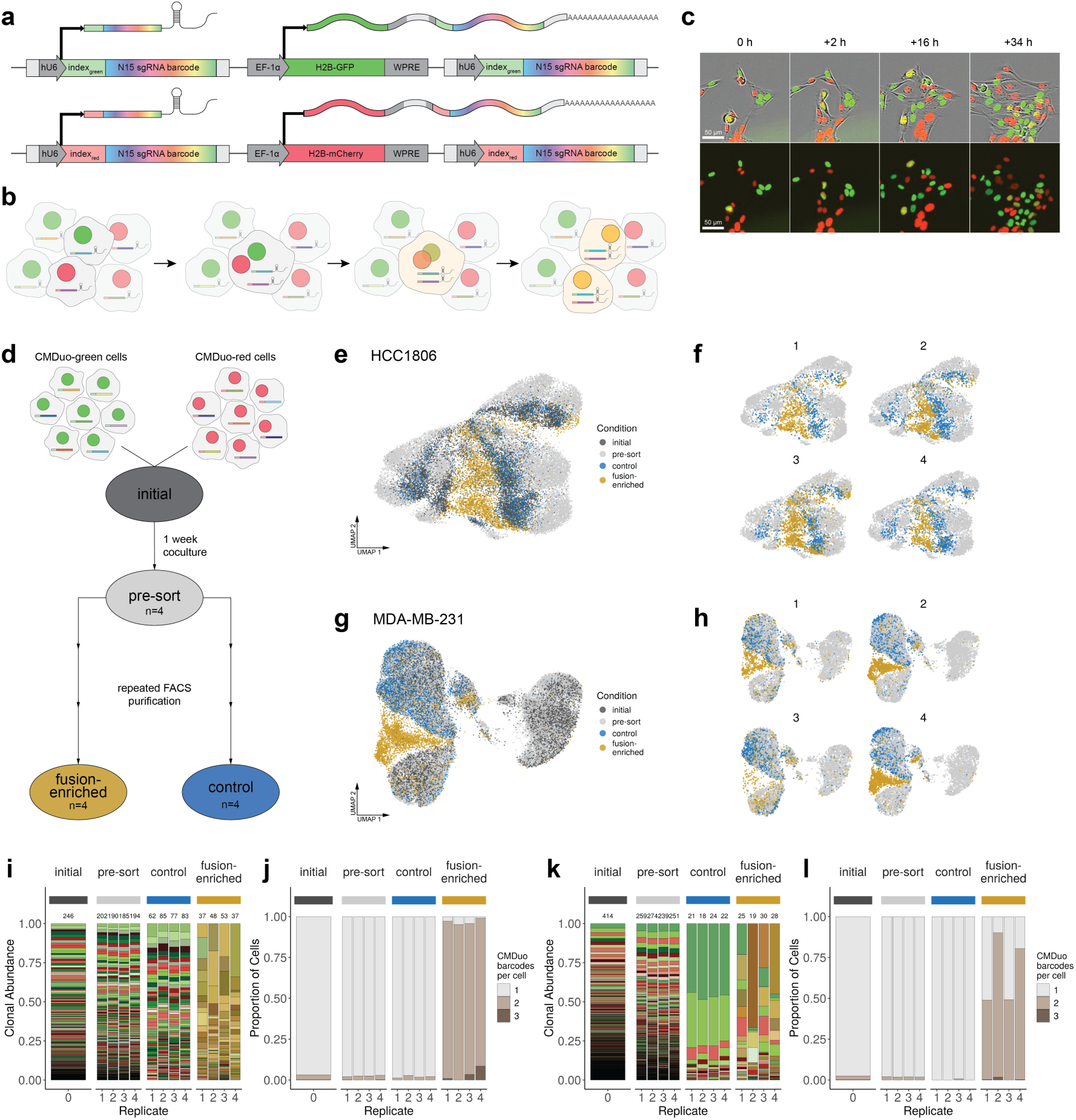
ClonMapper Duo is a two-color, two-index expressed DNA barcoding platform which enables tracking of cell-cell fusion events in scRNA-seq and live-cell imaging. (a) Schematic showing the core features of the CMDuo system. Cells with the integrated transgene express a barcode composed of a degenerate N(15) sequence with a 5-nt index sequence corresponding to the expressed fluorescent protein. The vector expresses the barcode as both a CRISPR-compatible sgRNA and a polyadenylated transcript. (b) Schematic following barcoded cells through cell-cell fusion, genomic recombination and division. Cells generated from cell-cell fusion of CMDuo-green and CMDuo-red cells will express barcodes and barcode-associated fluorescent proteins from both parental cells. (c) 10X live-cell imaging capture of a spontaneous cell-cell fusion event between CMDuo-red and CMDuo-green HCC1806 cells (additional images in Extended Data. Fig. 1). (d) 1:1 mixtures of homotypic CMDuo-green and CMDuo-red barcoded HCC1806 or MDA-MB-231 cells were cocultured in 5 independent replicates. One replicate was analyzed by scRNA-seq (“initial”) 2 days after plating. The remaining 4 replicates stayed in coculture for 1 week (“pre-sort”) before undergoing multiple rounds of FACS sorting to enrich for double-positive fusion events (“fusion-enriched”) or single positive (“control”) cells from each replicate for each cell line. (e) UMAP representation of scRNA-seq for all cells in the HCC1806 data set colored by sample and (f) shown for each replicate. (g) UMAP representation of scRNA-seq for all cells in the MDA-MB-231 data set colored by sample and (h) shown for each replicate. (i) Distribution of clones and (j) the number of barcodes assigned to each cell within each sample for HCC1806. Here a unique green, red, or gold/brown was assigned to each CMDuo-green, CMDuo-red, or fusion clone, respectively. (k) Distribution of clones and (l) the number of barcodes assigned to each cell within each sample for MDA-MB-231.

### ClonMapper Duo allows clonal population tracking and confident assignment of multi-barcoded fusion cells

To track spontaneous homotypic cell-cell fusion from parental clones to progeny in triple negative breast cancer cells, the CMDuo platform was applied as in Fig. 1d for two cell lines (HCC1806, MDA-MB-231) with four parallel replicates each. Homotypic cocultures were created by seeding an equal proportion of CMDuo-green and CMDuo-red barcoded cell libraries across replicate plates (“initial”). 1 week after coculture and expansion (“presort”), cells in each replicate were FACS sorted into two subsets double-positive (GFP^+^ mCherry^+^) fusion-enriched cells and single-positive (GFP^+^ or mCherry^+^) control cells. As no surface markers of fusion events are known to assist in discriminating fusion events from red-green doublets, multiple rounds of FACS were needed to obtain high purity fusion samples. FACS enrichment for fusion and control cells was repeated until fusion-enriched populations were > 50% pure. From fixation of the “initial” cells to the time at which “fusion-enriched” and parallel “control” cells were fixed for scRNA-seq, HCC1806 cells underwent 3 enrichment sorts over 35 days and MDA-MB-231 underwent 4 enrichment sorts over 50 days to achieve the final “fusion-enriched” and “control” samples for scRNA-seq analysis with the purity of each population summarized in Supp. Table 2.

In the initial population of HCC1806 cells, 246 unique clones (136 green and 110 red) were identified with a Shannon diversity index of 5.1 (evenness=0.92). This diversity drops minimally among initial, pre-sort, and control samples in the HCC1806 cells, with fewer barcodes and lower diversity observed in the fusion-enriched samples (Fig. 1i,k, Supp. Table 3). In the initial MDA-MB-231 cell library, 414 unique clones (197 green, 216 red, and 1 whose index sequence could not be confidently assigned) were identified with a Shannon diversity index of 5.4 (evenness=0.90). Unlike the HCC1806 cells, the MDA-MB-231 cells showed a strong loss of diversity in the control cells which had undergone repeated rounds of FACS in parallel with the fusion cells (Fig. 1i,k). The loss of diversity in the MDA-MB-231 cells may be due to clonal growth rate differences over the longer experimental timeline or stronger subclonal responses to repeated cell sorting. The number of barcodes assigned to each cell within each sample was quantified (Fig. 1j,l) and for all samples, the proportion of cells assigned multiple barcodes by scRNA-seq is well-matched to the proportion of fusion cells quantified via flow cytometry (Supp. Table 4, Supp. Fig. 2)

### High-resolution mapping of cell-cell fusion events

To map parental clones to fusion progeny, cells in the fusion-enriched sample containing both red and green barcodes were assigned as confident fusion events when at least 5 clonal cells containing the same barcode set were found in a single replicate. Fusion events will be annotated as outlined in Fig. 2a. 39.4% (97/246) of clones participated in at least one detected fusion event in the HCC1806 population, and 7.7% (32/414) in the MDA-MB-231 population (Fig. 2b-f, shown by replicate for both populations in Extended Data. Fig. 2). To test whether the clones found in the fusion population occur more often than expected by random chance, a binomial test was conducted, weighting the probability of selection by abundance in the initial population. 5.3% (13/246) and 2.1% (9/414) of clones in HCC1806 and MDA-MB-231 appeared more often than expected by random chance in the fusion clones (Fig. 2g-h). These results reject the null hypothesis of random selection and suggest that certain clones are more fusogenic or, if fusion is occurring randomly, that certain clones have innate features which better prime progeny for survival after fusion.

**Figure 2.**
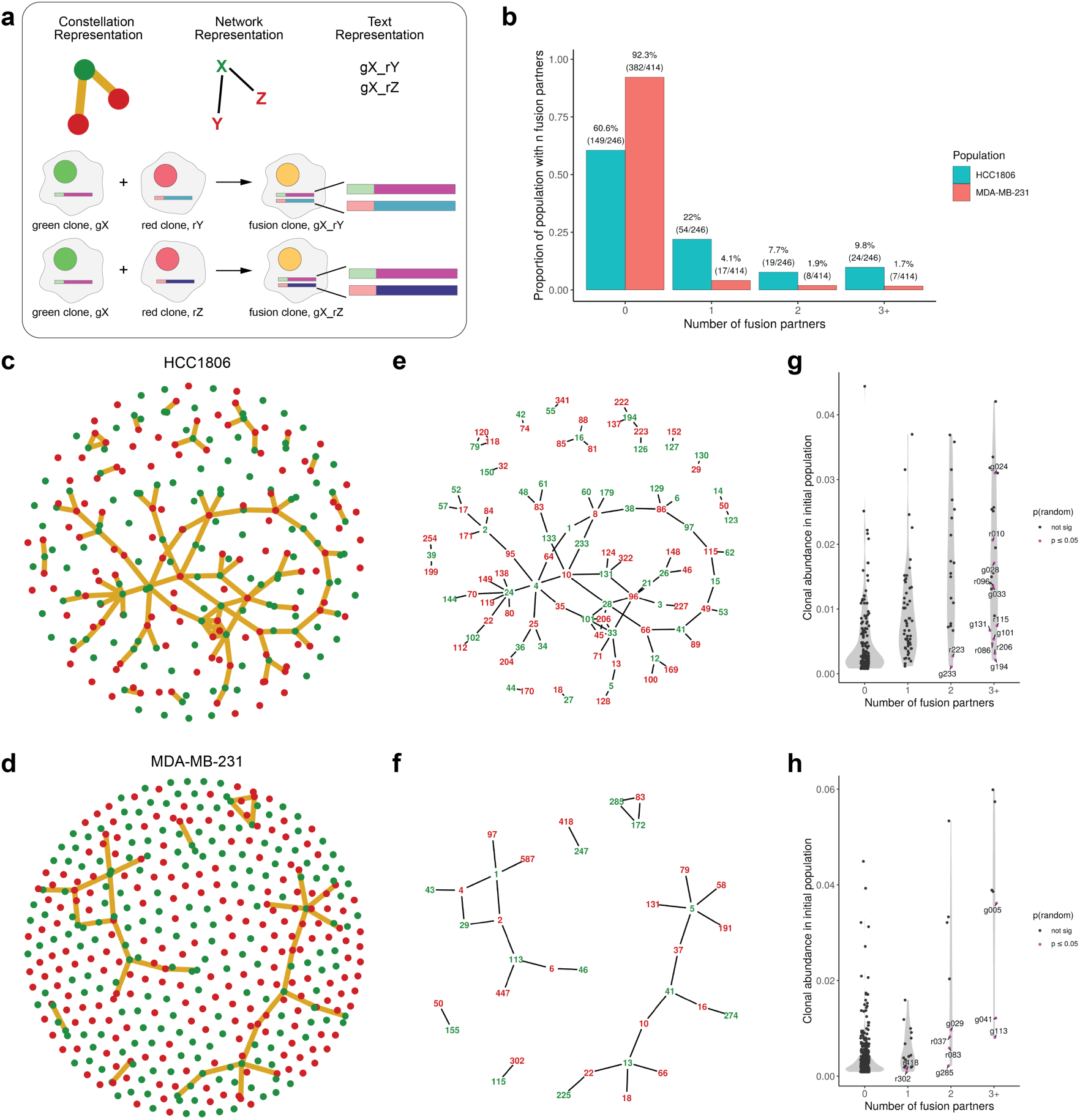
High-resolution mapping of cell-cell fusion events. (a) Legend showing an overview of different representations of fusion events, here we show how a fusion event between green clone ‘gX’ and two different red clones ‘rY’ and ‘rZ’ can be represented as constellation plots, network plots, and in text. (b) Proportion of clones participating in fusion events in each cell line. (c) Constellation plot showing all clones with edges representing a fusion event between clones in the HCC1806 and (d) MDA-MB-231 populations. (e) Network representation highlighting the barcode identifiers for clones involved in cell-cell fusion events for HCC1806 and (f) MDA-MB-231. (g) Quantification of the number of fusion partners for each clone, highlighting clones which participated in fusion events more often than random chance given abundance in the initial population for the HCC1806 and (h) MDA-MB-231 populations.

### Parental clones are enriched for actin and actomyosin-related pathways

To investigate whether there are transcriptomic features of parental clones which predict their occurrence in fusion progeny, parental clones which appeared more often in fusion progeny than expected by random chance (Fig. 2g-h, labeled points) to clones which were not found in any fusion progeny for each cell line using a mixed-model analysis. At a log2FC cutoff of 0.2 and adjusted p-value (p-adj) <= 0.05, there were 224 upregulated and 324 downregulated genes identified in HCC1806 and 2 up and 1 down in MDA-MB-231. Considering that the minimal number of significant genes in the MDA-MB-231 sample may be driven by lower sample size due to the lower rate and/or capture of fusion events in this cell line, we asked if any of the significant genes from the HCC1806 overlap with non-significant genes with a log2FC cutoff of 0.2 in MDA-MB-231 and find 37 potential markers (Supp. Table 5, Extended Data Fig. 3a-b). Although no significant genes were shared between cell lines, gene set enrichment analysis identified enrichment for actin and actomyosin-related pathways in both cell lines (Extended Data. Fig. 3c-d).

### Fusion progeny hybridize parental transcriptomic states and activate new programs

Next, we investigated the transcriptomic fate of fusion progeny by first asking if fusion clone clones maintain clone-specific transcriptomic patterns from their parental clones. Marker genes for parental clones in each parent-parent-progeny set were calculated and then their expression was analyzed for all clones within the matched parent-parent-progeny set, finding that fusion clones hybridize marker genes from their parental clones in all examples (Fig. 3a-i, Extended Data Fig. 4-5). To quantify transcriptomic similarity, the Pearson correlation coefficient was calculated for each parent-parent and parent-fusion pair in each parent-parent-fusion set. This revealed that parental clones in both populations are more similar to their fusion progeny than they are to each other (for HCC1806 (n=27) PCC_parent-parent_ = 0.87, 95% CI [0.82, 0.91], PCC_parent-fusion_ = 0.91, 95% CI [0.87, 0.91], and for MDA-MB-231 (n=12), PCC_parent-parent_ = 0.88, 95% CI [0.85, 0.95], PCC_parent-fusion_ = 0.92, 95% CI [0.88, 0.95] (Extended Data. Fig. 6). These results show fusion progeny inherit and hybridize clone-specific transcriptomic patterns from their parental clones.

**Figure 3.**
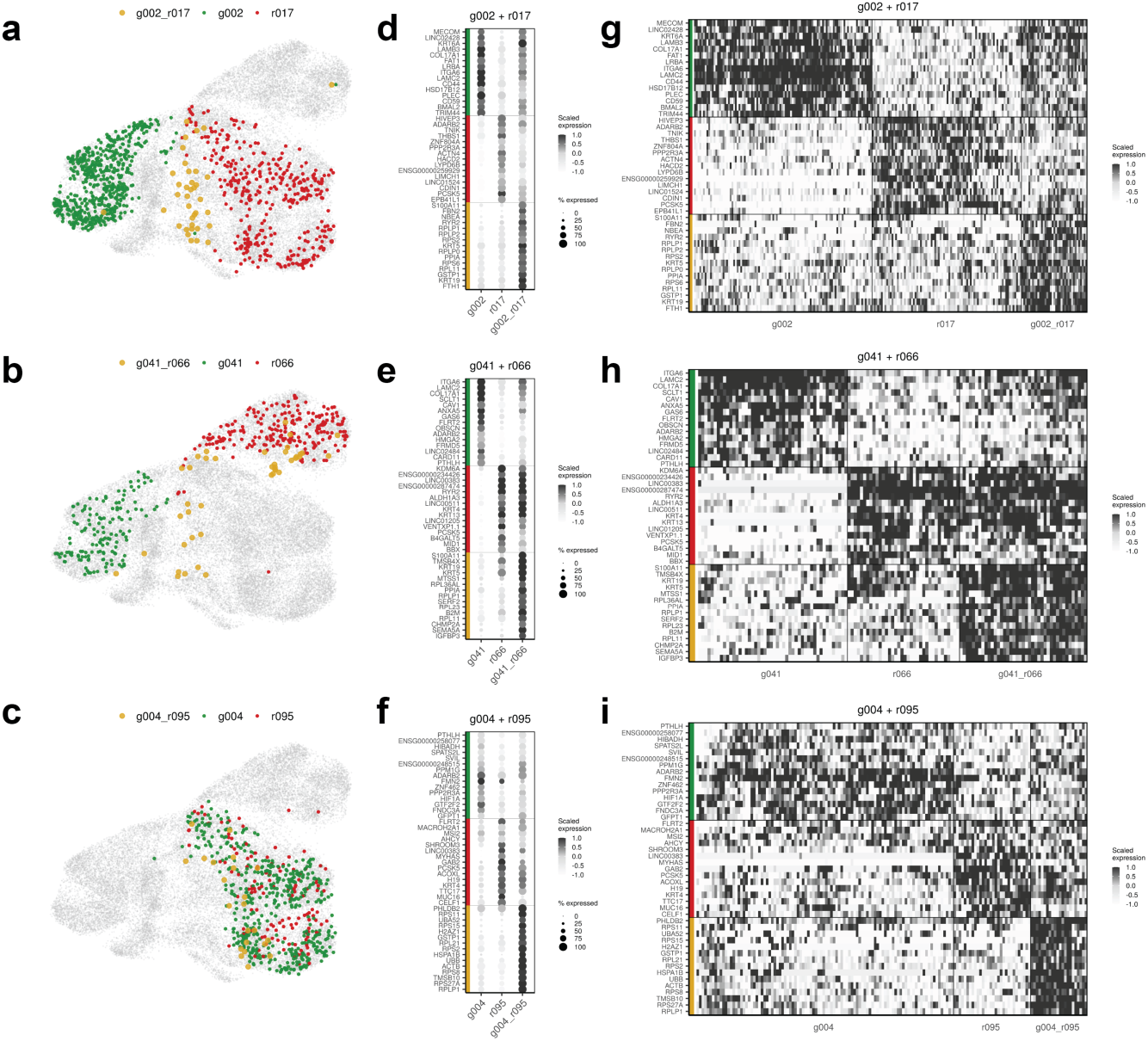
Fusion progeny hybridize parental transcriptomic states and activate new programs. Clone-to-clone analysis highlighting examples of 3 distinct fusion examples in the HCC1806 population: (top) g002+g017, (center) g041+r066, and (bottom) g004+r095. Showing for each (a-c) UMAP representations of each parent-parent-fusion set (d-f) average scaled expression of clone-specific marker genes for each clone in the parent-parent-fusion set where the intensity of the dot represents scaled expression and the size of the dot represents the number of cells of that clone expressing that gene, (g-i) scaled expression of clone-specific marker genes for each clone visualized across cells from each clone.

To investigate if novel cell states are activated after cell-cell fusion, differential expression analysis was performed between for each fusion clone against both of its parental clones. Here, in each fusion clone there is a set of genes that become highly active in fusion clones compared to the parental clones. This suggests that in addition to the hybridization of parental genes, the process of cell-cell fusion generates novel cell states.

### There is a distinct cell state achieved through cell-cell fusion independent of parental clone transcriptomic identity

Finding that fusion progeny not only hybridize parental cell states, but also activate distinct transcriptomic states at clonal resolution, we next sought to characterize general transcriptomic features of fusion progeny across the population. Cells were clustered on gene expression using the Louvain algorithm and clusters sharing highly similar distributions of barcodes were collapsed (Supp. Fig. 3). This resulted in 7 distinct barcode-informed clusters for the HCC1806 population (2 of 8 Louvain clusters collapsed), and 7 for the MDA-MB-231 population (0 of 7 Louvain clusters collapsed) (Fig. 4a-d, Extended Data. Fig. 7a-d). Marker genes were calculated for each cluster (Fig. 4i, Extended Data Fig. 7i) and are listed in Supp. Tables 6-7.

**Figure 4.**
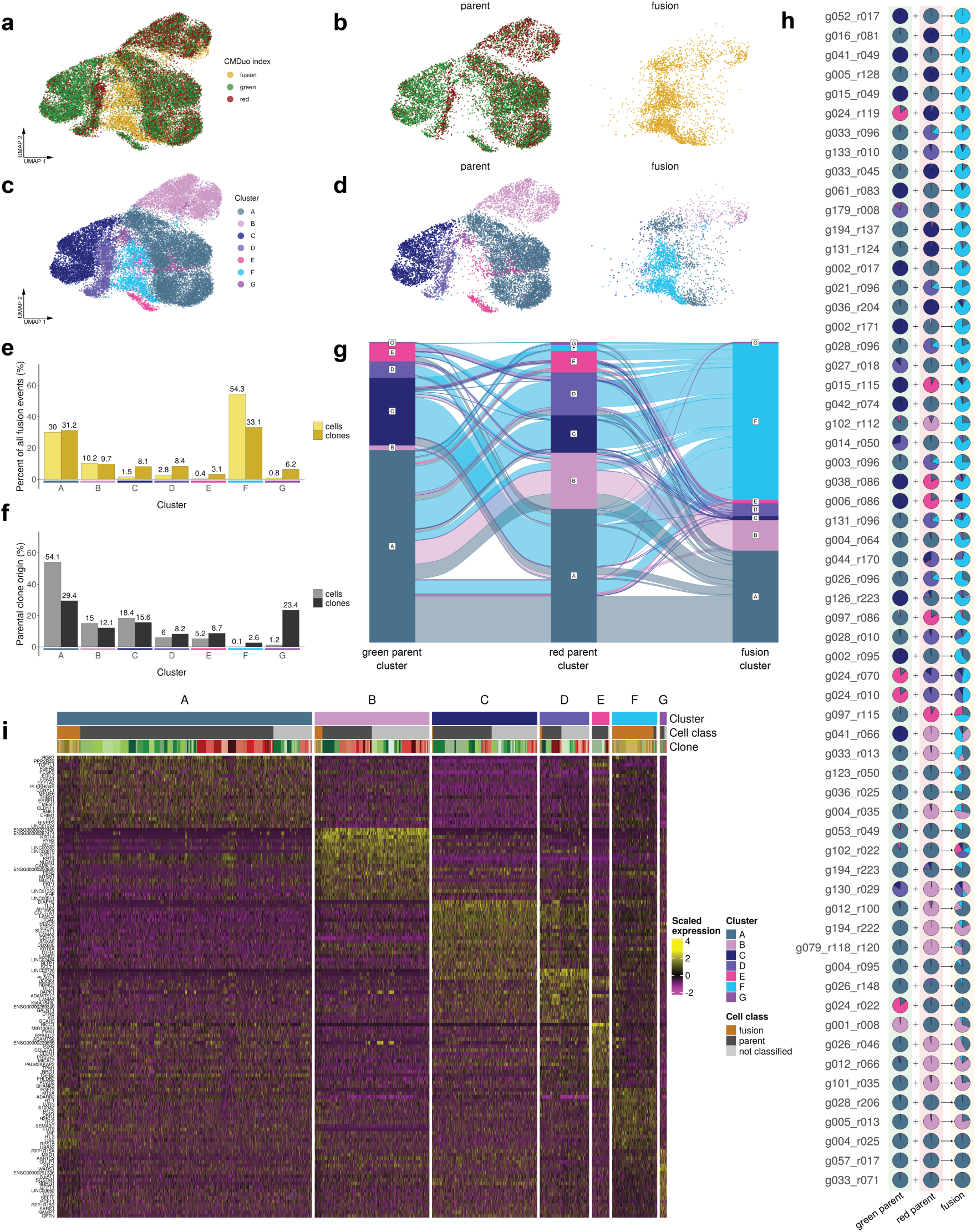
CMDuo reveals a distinct transcriptomic state driven by cell-cell fusion. UMAP representations of all barcoded cells colored by CMDuo index (a-b) or (c-d) transcriptomic cluster in the HCC1806 population (see Extended Data Fig. 7 for parallel analysis on MDA-MB-231). (e) Percent of all confident fusion events in each transcriptomic cluster showing percent of all fusion cells (light) and percent of all fusion clones (dark). (f) Origin of parental cells in each transcriptomic cluster showing percent of parental cells (light) and percent of parental clones (dark). (g) Alluvial plot summarizing the relationships between parental transcriptomic clusters and fusion progeny transcriptomic cluster. (h) Pie chart representation of the proportion of each clone in each cluster for each set of parent-parent-fusion clones. (i) Heatmap showing marker gene expression for each cluster.

For each cell line, fusion clones were found in all clusters present in the parallel sorted controls (Fig. 4a-e, Extended Data Fig. 7a-e) but the highest number of cells and unique clones was identified in Cluster F (54.3% of fusion cells and 33.1% of unique fusion clones in HCC1806 and 64.3% of fusion cells and 28.6% of unique fusion clones in MDA-MB-231) (Fig. 4e, Extended Data Fig. 7e). Parental cells were nearly absent in cluster F (0.1% of cells in both cell lines) (Fig. 3f, Extended Data Fig. 7f).

Given the distribution of fusion clones across different transcriptomic clusters, we asked if these different cell states were deterministically generated as a function of the transcriptomic identities of parental clones. We find that the parental transcriptomic cluster identity can be passed to fusion progeny if fusion occurs between clones from the same cluster, but the inverse is not true, there is a transcriptomic cluster that cannot be confidently mapped back to parental clone state. While the cluster F cell state is most often achieved through cluster-mismatched clones, it can also be achieved through fusion of any combination of parental cell states within the population (Fig. 4g-h, Extended Data Fig. 7g-h).

To determine features that distinguish the fusion-enriched cluster F cell state from other cell states, we identified overlapping differentially expressed genes and enriched gene sets across both cell lines. 466 differentially expressed genes (206 up, 260 down) at abs(log2FC) > 0.2 and p-adj <= 0.05 were identified in cluster F in both cell lines (Extended Data 8a) (genes at abs(log2FC) > 0.35 shown in Table 1. all shared significant genes in Supp. Tables 8-9), showing enrichment for pathways involved in chromatin remodeling, DNA replication/repair, and downregulation of pathways involved in activation of immune response (Extended Data Fig. 8c,e). This significant decrease in interferon signaling and immune signaling pathways in cluster F cells may indicate a key mechanism of survival for fusion progeny within tumor environments.

**Table 1.**
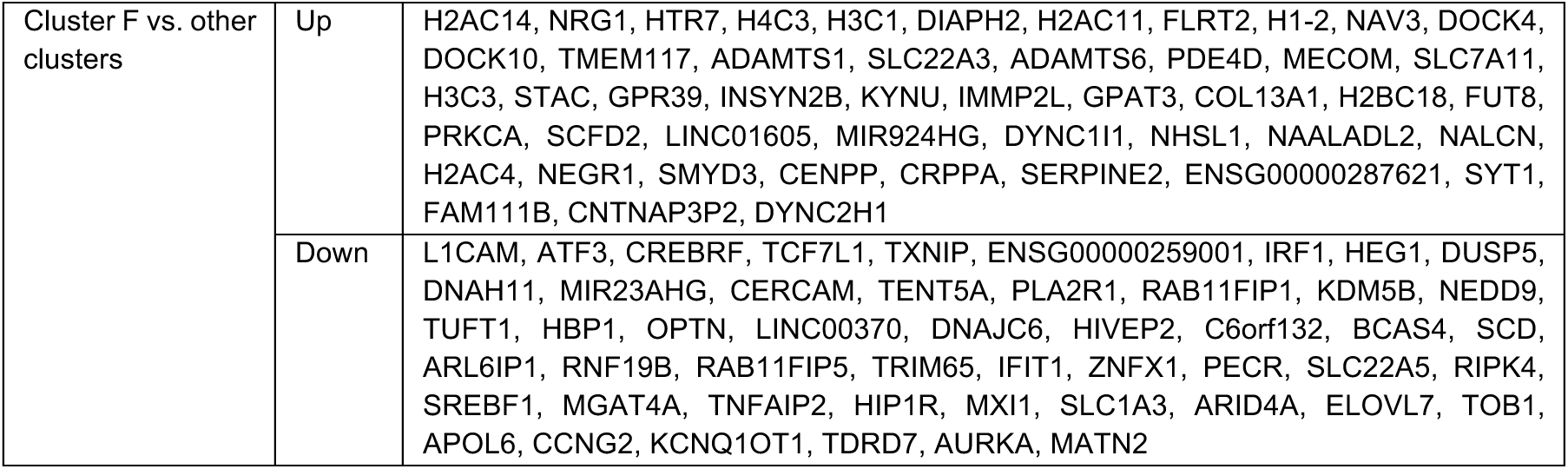
Top differentially expressed genes in Cluster F in both cell lines (abs(log2FC) > 0.35, p-adj <=0.05).

### Fusion progeny within parental clusters have transcriptomic signatures distinct from cluster F fusion cells

While the highest number of fusion cells were found in the cluster F, fusion cells in both cell lines were identified in other transcriptomic clusters. To first identify gene expression patterns altered in fusion clones across all clusters, fusion cells were compared to parallel control cells within each cell line. 117 differentially expressed genes were found (78 up, 39 down) (significant genes at abs(log2FC) > 0.35 shown in Table 2, all significant genes in Supp. Tables 10-11), and there is a pattern of an enrichment for E2F targets, reactive oxidative species, oxidative phosphorylation, and DNA repair across all fusion cells (Extended Data Fig. 9).

**Table 2.**
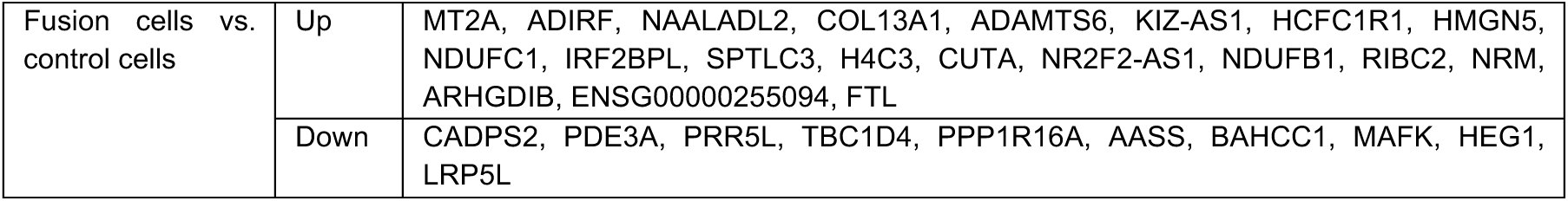
Top differentially expressed genes in fusion cells versus parallel control cells across all clusters (abs(log2FC) > 0.35, p-adj <=0.05).

Given that fusion clones could achieve multiple cell states by maintaining a state similar to their parental identity or activating cluster F programs (Fig. 4h, Extended Data Fig. 7h, Supp. Figs. 4-5), we next asked if fusion cells in cluster F are significantly different from fusion cells in other clusters and revealed striking differences. 585 differentially expressed genes shared in both cell lines (246 up, 339 down) were identified between fusion cells in cluster F and fusion cells in parental clusters (Fig. 5a-b) (abs(log2FC) > 0.2, p-adj <=0.05) (significant genes at abs(log2FC) > 0.5 shown in Table 3). Many of the top shared genes in cluster F fusion cells are histone genes, suggesting ongoing chromatin remodeling, supported by strong enrichment for nucleosome organization by GSEA (Fig. 5d). Compared to cluster F fusion cells, fusion cells found in their parental clusters show enrichment for proteotoxic stress and viral response related pathways (Fig. 5d). These fusion cells also show higher expression of mitotic genes (CCNB1, AURKA, PLK1, NEK2, KIF20A, CKAP2, CDC20, DEPDC1, CKS2, SGO2) and a higher proportion of cells in G2/M compared to cluster F (Extended Data Fig. 10g-h), which may suggest a more proliferative cell state.

**Table 3.**
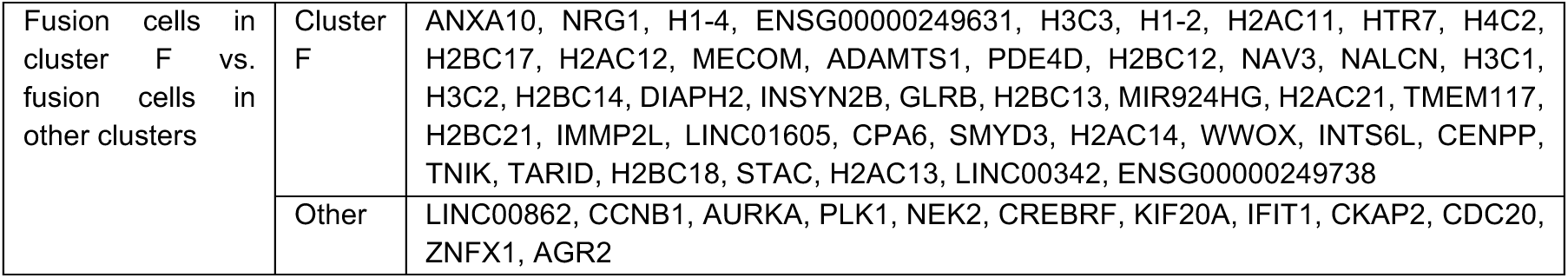
Top differentially expressed genes in fusion cells in cluster F vs fusion cells in other clusters (abs(log2FC) > 0.5, p-adj <=0.05).

**Figure 5.**
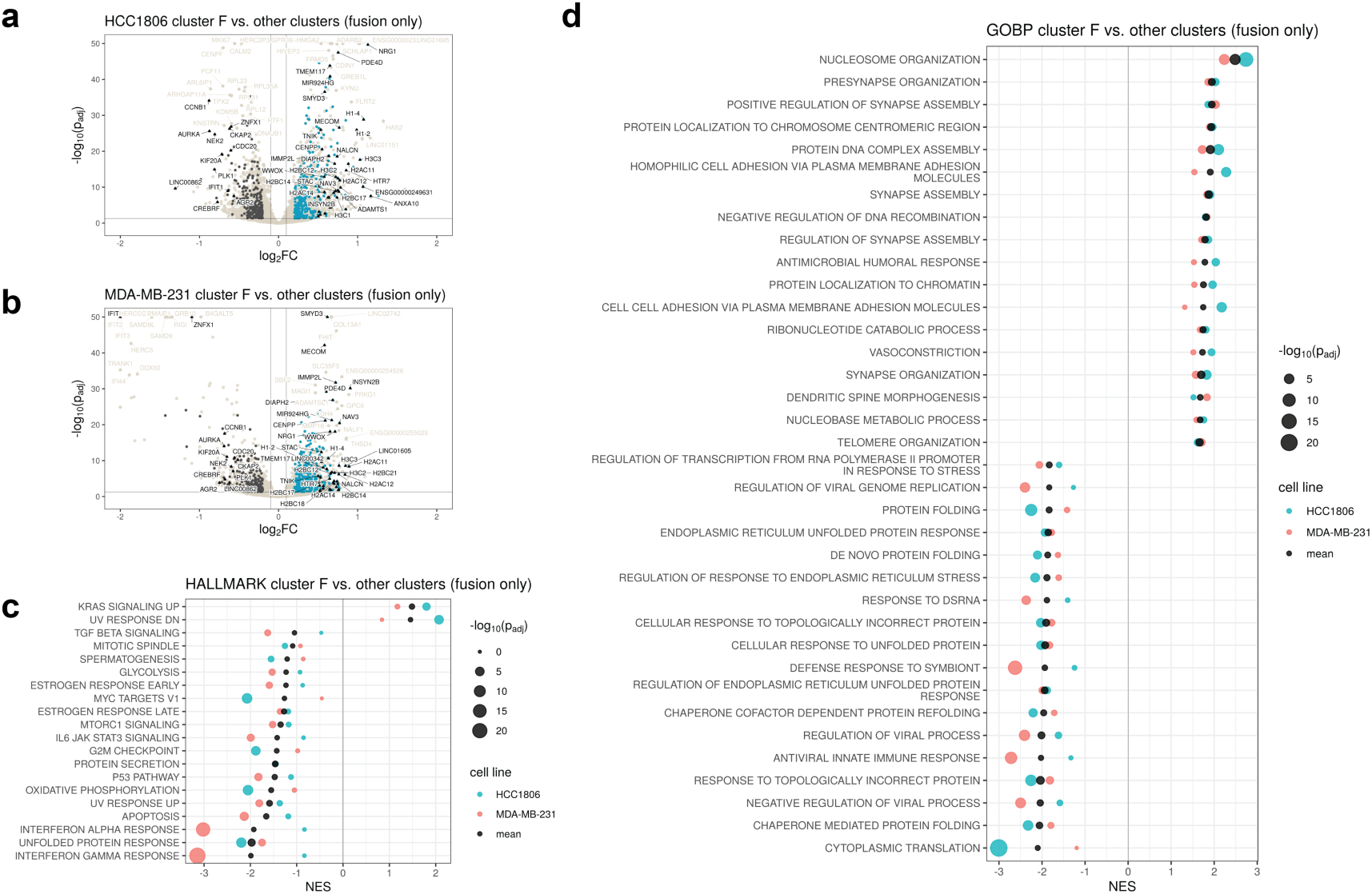
Differential expression and gene set enrichment analysis of fusion cells in cluster F versus fusion cells in other clusters. (a) Volcano plot of differentially expressed genes in cluster F fusion cells (blue, right) vs. other cluster fusion cells (dark grey, left) for the HCC1806 and (b) MDA-MB-231 populations, highlighting the top genes differentially expressed in both cell lines. (c) Hallmark enrichment analysis showing pathways up or down in both cell lines, with positive NES scores matched to cluster F fusion cells and negative NES scores for fusion cells in other clusters. (d) GO biological process enrichment analysis showing pathways up or down in both cell lines.

## Discussion

Cell-cell fusion in cancer confounds our current understanding of tumor evolution as an asexual process^14,32^, but it can no longer be ignored. Genome doubling through cell-cell fusion may be a mechanism by which cells are able to offset the negative effects of mutational load and slow Muller’s Ratchet^33,34^. There are well-studied cell types in the human body known to participate in fusion in normal and pathological contexts, but these events can be more readily classified due to maintenance of multiple nuclei. In this study and numerous others, multinucleation after cell-cell fusion was observed as a transient cell state producing single-nucleated fusion progeny visually similar to the parental cells (Fig. 1c, Extended Data Fig. 1). Without the use of red and green fluorescent markers, fusion cells may be indistinguishable from parental cells. This leads us to hypothesize that the rate of cell-cell fusion in cancer and in normal tissues may be higher than currently measurable. CMDuo now enables high-resolution *in vitro* investigation into the causes and consequence of cell-cell fusion in cancer and other tissues.

Prior studies investigating cancer cell-cell fusion have largely used heterotypic, often genotypically mismatched, and sometimes species mismatched cells^13,35–41^. Studies performed in these model systems have allowed assessment of parental contribution to the phenotypic and karyotypic state of fusion progeny, but many of the reported signatures have focused on inherited features from parental cell types. For example, it has been reported that fusion between immune and cancer cells produces tumor hybrid cells with leukocyte-specific pathways^23,42–44^, but the extent to which the physical process of cell-cell fusion may have further amplified or muted the acquired features and the role of subclonal phenotypes within each parental population has not yet been explored. With the resolution achievable using the CMDuo platform, we chose to investigate homotypic cell-cell fusion to decouple transcriptomic changes induced by the process of cell-cell fusion itself from changes driven by response to mismatched genotypes and states inherited from parental cells.

In the cell populations tested here, a small percentage of parent clones (Fig. 2b,g-h) with enrichment for actin and actomyosin-related pathways (Extended Data Fig. 3c-d) appeared more often in fusion progeny than expected by random chance. This may represent a phenotype which enhances survival and proliferation after fusion, or may suggest active actin restructuring toward a more fusogenic cell state^45,46^.

This study specifically enriched for and investigated cells which proliferate after cell-cell fusion, but cell-cell fusion does not always lead to a proliferative phenotype. Fusion progeny often do not survive fusion events^30^, they may remain multinucleated, or may adopt a giant polyaneuploid cell state. Multinucleated fusion cells and giant polyaneuploid cancer cells can become senescent or undergo eventual ploidy reduction to generate proliferative progeny^47–49^. In this timescale of this experiment, it is reasonable to assume that some of the fusion progeny collected and characterized at the final timepoint may have arisen from transition through a giant cell state. Further work is needed to quantify the occurrence of different cell-cell fusion outcomes in these populations and to link each outcome to parental cell states.

In this study, lineage tracing with ClonMapper Duo revealed that fusion progeny inherit and hybridize transcriptomic patterns from their parental clones and activate distinct transcriptomic programs in response to cell-cell fusion. We identified two distinct cell states achieved through homotypic cell-cell fusion often co-occurring within each fusion clone. The most common state achieved by cell-cell fusion was transcriptomically distinct and not correlated to parental cell state. This fusion-driven cell state was characterized by pathways related to chromatin remodeling, potentially suggesting a more epigenetically plastic cell state. Additionally, compared to other cells in the population, fusion progeny in the fusion-driven cell state had a marked decrease in interferon signaling suggesting a more immune privileged cell state which may have therapeutic implications. Fusion cells that adopted a transcriptomic signature more closely matched to their parental state were enriched for both oxidative phosphorylation and glycolysis and showed signatures correlated to an increased rate of mitosis. These fusion cells also showed enrichment for viral response pathways, which we hypothesize is due to presence of cytosolic DNA from micronuclei-mediated chromothripsis^50–52^.

The relationship between the two identified fusion cell states is currently unknown. Fusion cells may be bistable in both states or fusion cells may pass through the fusion-enriched cluster F state before stabilizing into a phenotype more closely matched to the initial population. Given the mitotic signatures in the fusion cells outside of cluster F, the latter hypothesis appears supported by work from Miroshnychenko et al. who showed an increase in the growth rate of fusion clones over passaging^13^. Longitudinal studies are needed to investigate the stability of these cell states and assess the degree of interconversion.

Application of the CMDuo platform in the context of this study has revealed novel insights into cancer cell-cell fusion, but it is not without shortcomings. To prevent gene signatures associated with response to antibiotics from skewing transcriptomic results, we chose to engineer the CMDuo system without antibiotic selection features. While we were able to achieve relatively pure fusion samples for all HCC1806 replicates and 2 of the 4 MDA-MB-231 replicates during the experiment (Supp. Table 2, Supp. Table 4), the cells needed to undergo multiple rounds of cell sorting and recovery to reach this level of purity. The inclusion of FACS controls in this study assisted with regressing out changes in gene expression due to cell sorting, but the repeated sorting process itself was a stronger selective pressure than expected in the MDA-MB-231 model. In a prior study, two highly distinct transcriptomic clusters were described within the MDA-MB-231 cell line, characterized by high or low expression of the surface marker, ESAM^53^. Here, the ESAM-low subpopulation was depleted in both the fusion and control samples (Supp. Fig. 6, Fig. 1h, Extended Data Fig. 7d) and the same 2 clones were selected for in control samples across all four replicates (Fig. 1k). There may be significant fusion occurring in the ESAM-low MDA-MB-231 subpopulation, but robust inferences cannot be made given the degree of FACS-induced selection seen in the control samples and the low number of fusion and control cells in this subpopulation. Antibiotic selection with the proper controls, may have been a more gentle and efficient method to enrich for double-barcoded fusion cells. Future iterations of the CMDuo system should include antibiotic selection as an optional feature.

ClonMapper Duo is a novel platform which was used to study homotypic cell-cell fusion in triple-negative breast cancer cells but can be used to study homotypic or heterotypic cell-cell fusion between any mammalian cell types *in vitro*. Additionally, this study highlighted the use of CMDuo for tracking cell-cell fusion, but it can be readily applied to lineage tracing studies with other goals. The fluorescence-associated barcodes allow confident assignment of population identity in scRNA-seq; different cell populations can be cocultured and accurately demultiplexed in scRNA-seq which will be useful for investigation of other cell-cell interactions. Further, the localization of the barcodes to the nucleus with H2B-GFP or H2B-mCherry allows precise counting of population size with live-cell imaging, which is useful for determining population growth rates and fitting data to mathematical models of cancer^54,55^.

Without biomarkers, the clinical relevance of cell-cell fusion in cancer remains unknown^35^. In this study, we applied a novel barcoding platform and unveiled signatures of clonal population before and after cell-cell fusion. Future studies into the mechanistic role of the identified gene signatures are required, and these studies may lead to the discovery of biomarkers for assessing fusogenic potential and the ability to quantify the prevalence of fusion in patient samples. If cell-cell fusion is identified as a key component of tumor evolution in any cancer type, then targeting pathways involved in fusogenic potential or response to cell-cell fusion may represent novel treatment strategies.

## Methods

### Construction of the CMDuo plasmids

The H2B-GFP gene block amplifying was cloned from the FU-H2B-GFP-IRES-Puro plasmid (AddGene #69550) by performing a 1X Q5 PCR reaction. Briefly 25 µL Q5 High-Fidelity 2X Master Mix, 2.5 µL of 10uM forward primer, 2.5 µL of 10uM reverse primer, 0.5 ng of template DNA, and nuclease-free water to 50 uL were mixed then PCR amplified with the following settings: (1) 98C for 2 min, (2) 98 °C for 5 sec, (3) 72 °C for 30 sec, (4) 72 °C for 1 min, (5) Repeating steps 2-4 for 20 cycles, then (6) 72 °C for 2 min, and (7) 4 °C hold. The PCR product was isolated and purified using gel electrophoresis. H2B-mCherry was ordered as a gene block designed to match the resulting H2B-GFP gene block but with mCherry in place of GFP. To linearize and remove the BFP sequence from the ClonMapper vector backbone (CROPseq-BFP-WPRE-TS-hU6-BsmBI, AddGene # 137993), 3 ug of plasmid was digested in a 3X restriction enzyme reaction containing 15 uL Digestion rCutSmart Buffer (NEB, B6004), 3 µL BsiWI-HF (NEB, R3553), 3 µL MluI-HF (NEB, R3198), and nuclease-free water to 150 µL at 37 °C for 1 hour. The linearized plasmid band was purified using gel electrophoresis. Gene blocks were then separately ligated into the linearized ClonMapper backbone at a molar ratio of 2:1 in a 1X HiFi Assembly reaction by mixing 0.0638 pmol PCR product, 0.0319 pmol linearized plasmid, 10 µL HiFi DNA Assembly Master Mix (NEB, E2621), and nuclease-free water to 20 µL at 50 °C for 1 hour. 2 µL of the finished reaction mixture was heat shocked into one tube of competent e. coli (NEB, C3040) at 42 °C for 30 seconds, incubated in recovery media for 1 hour, then spread over ampicillin agar plates for overnight incubation. Colonies were expanded and the insertion of the reporter sequence was confirmed with Sanger sequencing.

### Generation of indexed barcode plasmid libraries

Random indexed barcode library gene bocks for GFP and mCherry were made by mixing equimolar ratios of each of their respective 4 random barcode templates (Supp. Table 1) and then performing an extension reaction as previously published^56–58^. The random indexed barcode gene blocks were inserted into their respective plasmid backbones using a BsmBI-v2 golden gate reaction following the published procedure. Targeted barcode sequencing was performed on the generated plasmid barcode libraries sequencing to assess population diversity following the published procedure.

### Cell culture

HCC1806 cells were cultured in RPMI with 2.05 mM L-glutamine (HyClone, SH30027.01), supplemented with 10% FBS (Sigma, F0926-500mL). MDA-MB-231 cells were cultured in high glucose DMEM (Sigma D5796) supplemented with 10% FBS (Sigma, F0926-500mL) and 1X penicillin-streptomycin (ThermoFisher, 15140122). MDA-MB-231 cells were passaged with 0.05% Trypsin-EDTA (ThermoFisher, 25300062) for 5 minutes and HCC1806 cells were passaged with 0.25% Trypsin-EDTA (ThermoFisher, 25200056) for 12 minutes when cultures reached 70% confluence. MDA-MB-231 cells were obtained from ATCC and barcoded within 5 passages. HCC1806 cells in the lab were STR verified before use.

### Generation of barcoded cell libraries

Lentiviral production and low MOI transduction (MOI=0.01) was performed according to previously published methods for the ClonMapper system^56–58^. For each cell line, CMDuo-green and CMDuo-red populations were generated in parallel from the same starting population by sorting replicates of 500, 800, or 1000 cells for each cell line. Sorted cell libraries were expanded and targeted barcode sequencing was performed to assess number of unique clones and skew within each sorted population. Low cell number libraries showing the highest evenness were chosen for the experiments. The HCC1806 populations selected for the scRNA-seq experiment came from expanded cell libraries of 500 CMDuo-green and 500 CMDuo-red sorted cells, and the MDA-MB-231 populations came from 800 CMDuo-green and 800 CMDuo-red sorted cells.

### Cell sorting and sample collection

For each cell line, five replicate plates were seeded, containing 1e6 cells each. One plate was fixed for scRNA-seq 2 days after seeding (“initial”), the other 4 plates were expanded in culture as independent replicates for one week before the first FACS enrichment for fusion cells. At the time of sorting, 0.5e6 cells were fixed for Parse scRNA-seq fixation and 0.5e6 cells were pelleted for DNA and targeted barcode sequencing. These cells comprise the “pre-sort” samples. The remaining cells were FACS sorted for fusion cells (both GFP^+^ and mCherry^+^ cells) and control cells (single GFP^+^ or single mCherry^+^). As FACS is imperfect at double discrimination and there are no additional surface markers currently known which can be added in to discriminate fusion events from red-green doublets, multiple rounds of FACS were needed to obtain high purity fusion samples. Given that multiple rounds of FACS may act as a selective pressure and cause transcriptomic shifts, control cells (GFP^+^ or mCherry^+^) were sorted in parallel with fusion cells (GFP^+^mCherry^+^) to correct for any selective pressures and transcriptomic shifts which may be driven by cell sorting stress rather than fusion in downstream analysis. Pre-sort purity and cell yield for each round of FACS is summarized in Supp. Table 2. Sorted cells were allowed to recover from cell sorting and expanded before undergoing repeat rounds of cell sorting until fusion populations were > 50% pure, then “fusion-enriched” and “control” cells were prepped for scRNA-seq and DNA extraction. From fixation of the “initial” timepoint to the time at which “fusion-enriched” and parallel “control” cells were fixed for scRNA-seq, HCC1806 cells underwent 3 enrichment sorts over 35 days and MDA-MB-231 underwent 4 enrichment sorts over 50 days. The final purity of each sample fixed for scRNA-seq assessed by flow cytometry is summarized in Supp. Tables 2, 4.

### Bulk targeted barcode library preparation

DNA was extracted from frozen cell pellets using the Invitrogen PureLink Genomic DNA kit (ThermoFisher, K182001) according to manufacturer’s instructions. Amplification steps were performed following published ClonMapper barcode amplification procedures^56–58^. Briefly, DNA was quantified with NanoDrop, then 1 µg of DNA was loaded into the first PCR reaction. Amplified DNA was cleaned with SPRI beads, then 4 ng of cleaned stage 1 product was loaded into the next PCR reaction to add unique Illumina i7/i5 indices to each sample. Stage 2 samples were cleaned with SPRI beads, then quantified Qubit using the 1X dsDNA high-sensitivity assay (ThermoFisher, Q33231). Illumina indexed samples were run on a NovaSeq 6000 (S1 flow cell, single read, 100 cycles) by the UT Genome Analysis and Sequencing Facility, targeting 100e6 reads for all samples and biasing more reads toward the earlier timepoint samples. ∼14e6 reads were obtained for each initial population, ∼6e6 for each pre-sort sample, and ∼3e6 for each control and fusion sample in each cell line.

### Bulk targeted barcode sequencing analysis, barcode naming convention, and creating a reference list for Parse CRISPR Detect alignment

The results of targeted barcode sequencing performed on bulk DNA extracted from each sample were combined into early and late timepoint groups for each cell line, with the early group containing the initial and pre-sort samples, and the fusion-enriched and control samples in the late group. The set of barcodes in each group was determined with pycashier^59^ (version 2024.1001) (https://github.com/brocklab/pycashier/), setting the Levenshtein distance to 2 and the clustering ratio to 1. Post-processing on the pycashier outputs was performed to remove noisy sequences detected in both cell lines and ensure a minimum Levenshtein distance of 2 between all pairs of barcodes within each cell line. If any similar barcodes were detected, they were collapsed into the more abundant sequence. Barcode color assignment was performed by matching the first 5 nucleotides of each detected sequence to the designed index sequences, (withGFP being one of the following sequences: GACAA, GCTGT, ATCGC, ACGCA, and mCherry being: CATCC, CGAGA, TAGTG, TCAAC) and allowing a Levenshtein distance of 1 to account for sequencing errors. Unique nucleotide barcodes were assigned names according to their fluorescence reporter and their relative abundance in the ranking abundance, i.e. g001 and r001 represent the most abundant CMDuo-green and CMDuo-red clones in the early timepoint samples via targeted sequencing. Named barcodes and sequences were formatted into a reference list for input to the Parse CRISPR Detect alignment pipeline (v 1.3.1), setting the barcode name as the ‘Guide_Name’, the detected sequence as both the ‘Guide_Sequence’ and ‘Target_Gene’, and including flanking regions around the barcode as the ‘Prefix’ (ATCTTGTGGAAAGGACGAAACACCG) and ‘Suffix’ (GTTTTAGAGCTAGAAATAGCAAGTT).

### Parse scRNA-seq library prep

Library prep was performed according to manufacturer guidelines outlined in CRISPR Detect for Evercode WT v2 (User Manual Version 1.1, UM0025). As the WT kit can process 100,000 cells, samples from both cell lines were prepared with the same kit. For each cell line, the ‘initial’ sample was split over 4 wells targeting 13,000 cells total, each of the four ‘pre-sort’ samples were split over 3 wells targeting 6,250 cells per sample, and each of the sorted control and fusion-enriched samples received its own well, targeting 1500 cells per sample. 8 identical sublibraries were prepared, but one sublibrary was lost during preparation, adjusting the expected cell number to 87,500 total and 11,375, 5,469, and 1312 per sample per cell line, respectively. ClonMapper Duo gRNA barcodes were amplified from sublibrary cDNA according to manufacture instructions for CRISPR Sequencing Library Preparation in CRISPR Detect for Evercode WT v2 (User Manual Version 1.1, UM0025). Whole transcriptome and CRISPR detect (CMDuo barcode) libraries were amplified separately from cDNA and sequenced independently across 8 sublibraries each. Whole transcriptome and CRISPR Detect sublibraries were sequenced at the UT Austin Genomic Sequencing and Analysis on a NovaSeq 6000 (S2 flow cell, paired-end, 200 cycles). 500e6 reads were targeted for each WT sublibrary and 50e6 reads for each CRIPSR Detect sublibrary, producing 41,973 mean reads per cell across transcriptomic libraries and 3,238 means reads per cell in the combined CRISPR Detect sublibraries.

### scRNA-seq genome annotation and sequence alignment

All sequence alignment and sample annotation steps were performed with Parse Bioscience pipeline (v 1.3.1) in a Linux environment running Python version 3.10. Genome annotation and reference files containing eGFP and mCherry sequences were concatenated to the most recent human genome build (GRCh38, release 111). A Parse reference genome was built from the concatenated genome and annotation files. Each Parse whole transcriptome sublibrary was aligned to the built genome, then aligned sublibraries were combined for analysis.

### scRNA-seq barcode alignment

Alignment and sample annotation steps were performed with Parse Bioscience pipeline (v 1.3.1) in a Linux environment running Python version 3.10. Formatted pycashier outputs from independently run targeted barcode sequencing were used as inputs to the Parse CRISPR Detect pipeline, setting a minimum read threshold of 3 and a minimum transcript threshold of 3.

### scRNA-seq quality control filtering and normalization

Quality control steps were performed on the unfiltered outputs of the Parse pipeline using Seurat (v5.1.0). Cells in retained for analysis had total RNA counts between 1,500-100,000 genes, total feature counts between 800-50,000 counts, and mitochondrial percent between 2-18%. Doublet detection was performed using scDblFinder^28^ (v1.18.0) and cells classified as doublets were removed. Cells were split by cell line and data from each cell line was independently normalized using a log transformation. Variable features were identified using a variance stabilizing transformation. Cell cycle score was calculated for each cell and scaled data from cell cycle regression was used to identify transcriptomic clusters and generate UMAP projections, but downstream analysis was carried out on non-cell cycle regressed data. Louvain clustering was performed using the first 20 principal components to identify transcriptomic neighbors and at a clustering resolution of 0.18 in both cell lines. 8 clusters were identified for HCC1806 and 7 clusters for MDA-MB-231 before barcode-informed clustering correction.

### scRNA-seq assignment of barcodes to cells

Barcodes were assigned to each cell using a probabilistic model accounting for sequencing noise in the data using the raw outputs from the Parse CRISPR detect pipeline. Assigned barcodes were first filtered to match the expected cell line, to have at least 3 reads per barcode in a cell, and to ensure that detected barcodes is present in at least 3 cells at this threshold within a sample. After this minimal filtering, the expected noise generated from each barcode was calculated based on overall barcode abundance in the population and the number of reads per cell. A negative binomial model was used to determine the probability that a barcode within a cell is noise. Barcodes with a p-adjusted value <= 0.05 were assigned to cells. Cells with multiple instances of green barcodes or multiple instances of red barcodes were indicated with underscores, e.g., gXXX_gYYY. Once barcodes were assigned, clone color was assigned as green, red, or fusion by presence of ‘g’ and/or ‘r’ in the assigned barcode name.

### scRNA-seq barcode-informed clustering correction

Given the that most clonal lineages maintain transcriptomic similarity^57^, we corrected for potential over clustering from the Louvain algorithm by quantifying clonal abundance within each cluster and combining Louvain clusters with similar clonal profiles. Distance between vectorized clonal abundance in each cluster was quantified using the Canberra method. Hierarchical clustering was performed to visualize similarity of barcode distributions within clusters and create a dendrogram tree. The tree was cut at a minimum distance of 200 in both cell lines. This resulted in no changes to the MDA-MB-231 population, retaining its 7 Louvain clusters, and one change to the HCC1806 population, collapsing 2 of the 8 clustering into one, and producing 7 barcode-informed clusters for downstream analysis in each cell line (Supp. Fig. 3).

### scRNA-seq cluster naming

Barcode-informed clusters were re-named from numeric values to letters A-G in each data set, setting cluster F in each to be the cluster with the highest quantity of fusion cells. Across data sets, cluster F is the only cluster which has any informed similarity. Cluster names A-E, and G were assigned to the remaining numerical clusters within each cell line without preference.

### Cell classification

Unique combination of at least one red and green barcode appearing in at least 5 cells within a fusion-enriched replicate were classified as “fusion” clones. The parental barcodes associated with each classified fusion clone were then used to classify cells as “parental” clones.

### scRNA-seq differential expression analysis

Cluster marker genes were identified using all cells in each data set using a Wilcoxon rank sum test, assigning the highest significant log2FC values as maker genes for each cluster. A mixed model approach with NEBULA^60^ was used for all other differential expression analyses to correct for underlying clonal composition, setting cell barcode as the random effect (i.e. the subject id). Unless otherwise stated, a minimum of 5 cells per classified clone was required to be included in each analysis. Output log fold change was converted to log2FC and p-values were adjusted using a Benjamini-Hochberg correction. The adjusted p-values, p-adjust, were used to determine significance. Differentially expressed genes were defined at an absolute log2FC threshold of 0.2 and a p-adjust value <= 0.05.

### Gene set enrichment analysis

Genes from NEBULA mixed model analysis were pre-ranked using log2FC and GSEA was performed using fgsea^61^ (v1.30.0) on the v2023.2 h.hallmarks and c5.go.biological_process gene sets from the MSigDB^62,63^. Gene sets displayed in plots show gene sets that were up or down in both cell lines and had a false discovery rate of less than or equal to 0.25 in at least one cell line.

### Parent-progeny marker gene analysis

For each parent-parent-fusion set with at least 20 cells in each fraction, parental marker genes were calculated using a Wilcoxon rank sum test between cells of the green parent clone and cells from the red parent clone. The top 15 differentially expressed genes for each were considered parental marker genes and used for visualizations in Fig. 3. Clonal similarity was calculated by pseudobulking gene expression for each clone, then calculating pairwise Pearson correlation coefficient between clones in each parent-parent-fusion set.

## Acknowledgements

We are grateful for funding from the NIH which supported this project: NIH F31CA268833 (to A.L.G.), R01CA226258, R01CA255536, and U01CA253540 (to A.B.) We thank the core facilities at UT Austin for access to their services and knowledge. Sequencing was performed by the Genomic Sequencing and Analysis Facility at UT Austin, Center for Biomedical Research Support (RRID: SCR_021713). Flow cytometry and FACS was performed at the Center for Biomedical Research Support Microscopy and Imaging Facility at UT Austin (RRID: SCR_021756).

## Data Availability

Raw and processed single cell RNA-sequencing can be downloaded from GEO with accession number GSE286213. Targeted barcode sequencing to generate a list of barcodes in the samples can be downloaded from Gene Expression Omnibus (GEO) with accession number GSE286212.

## Code Availability

Code used to process data and generate figures in this study can be found at www.github.com/brocklab/CMDuo-analysis

