## Extended Data Figures for "Mapping cell-cell fusion at single-cell resolution"

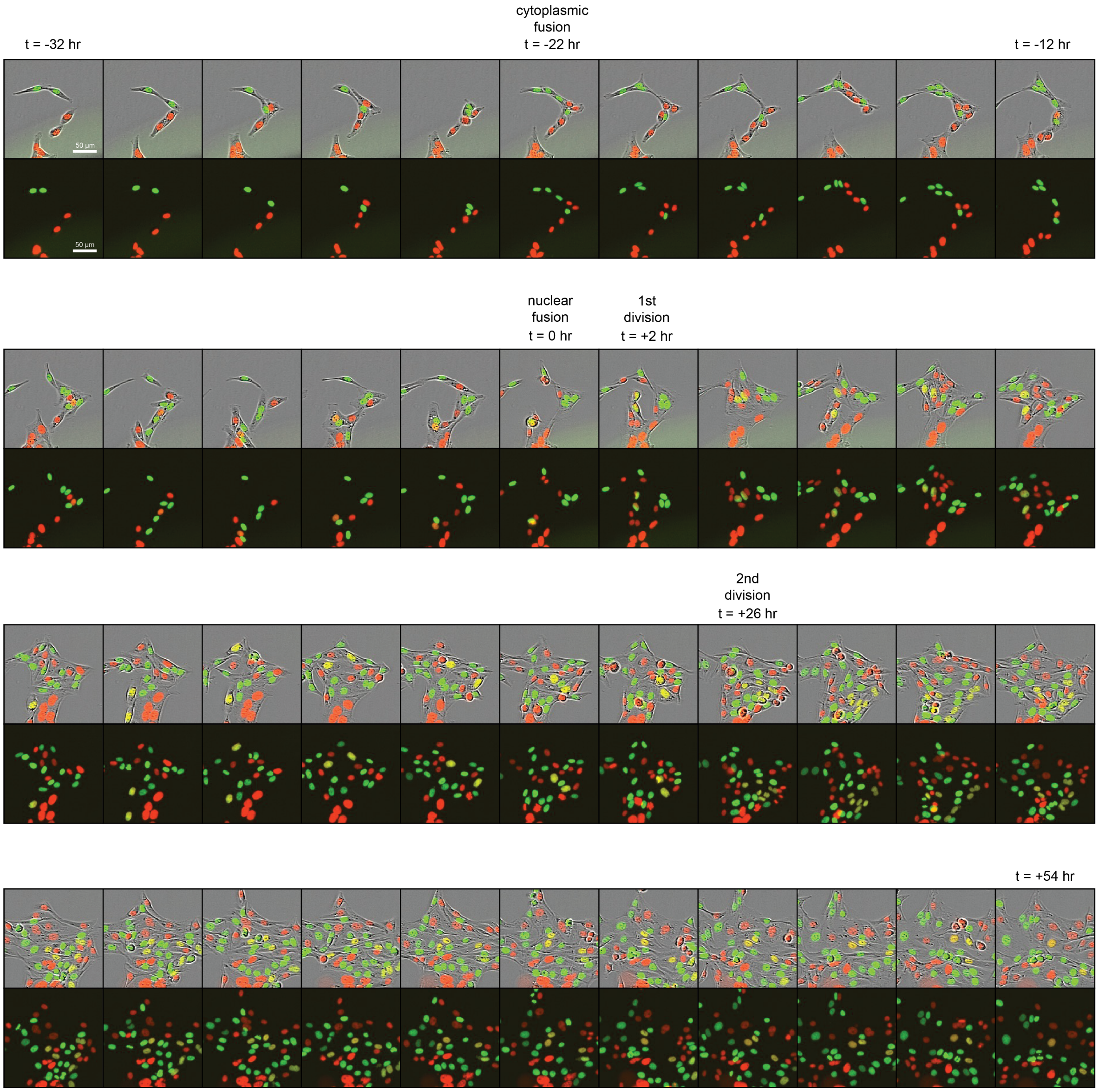


Extended Data Figure 1. Time lapse images of an HCC1806 fusion event occurring then passing through two rounds of cell division. Images were taken every 2 hours with a 10X objective in an IncuCyte S3. Scale bar is 50 µm.

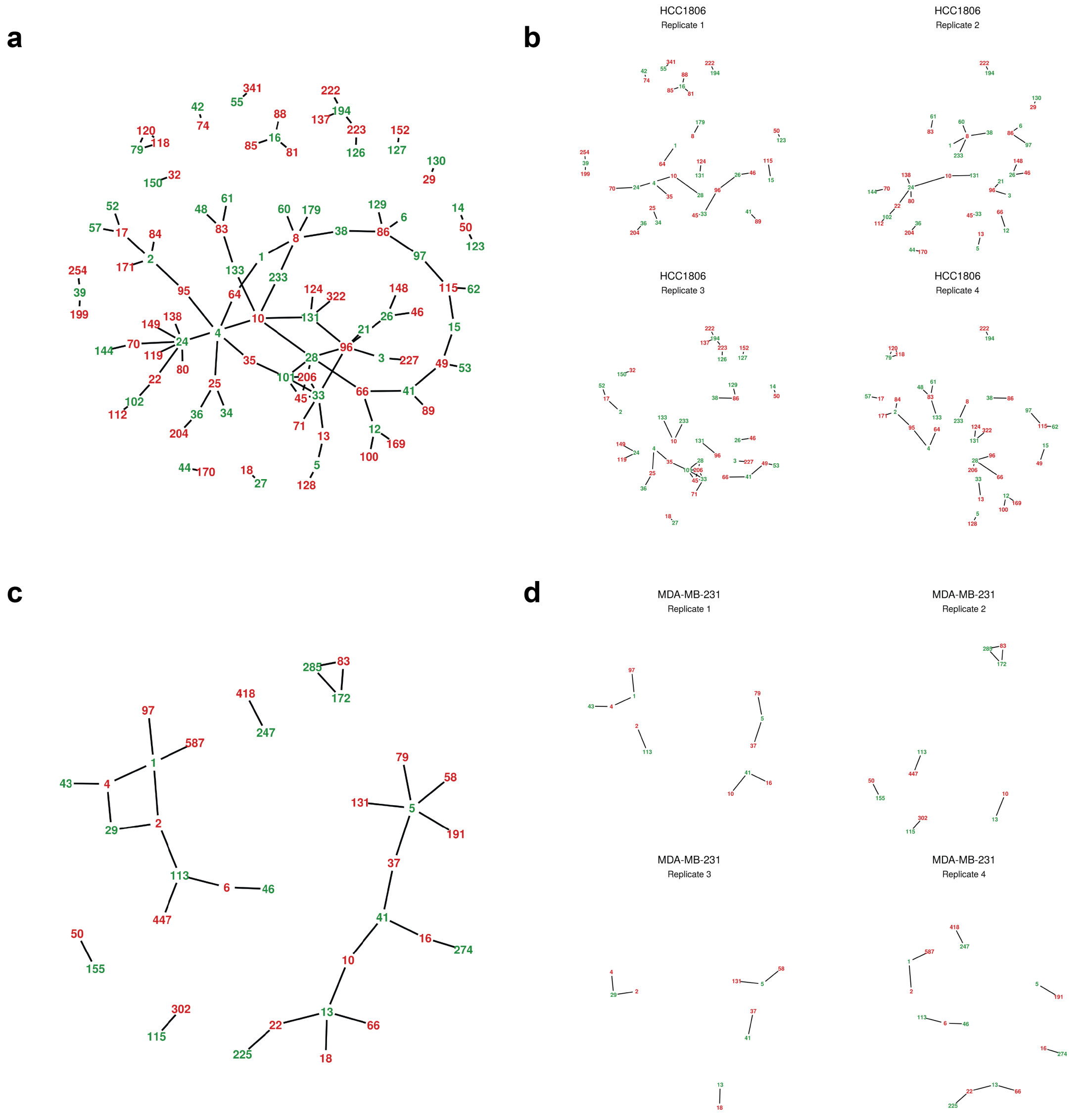


Extended Data Figure 2. Network plots showing all fusion events within the (a) HCC1806 and (c) MDA-MB-231 populations, and (b, d) showing each replicate.

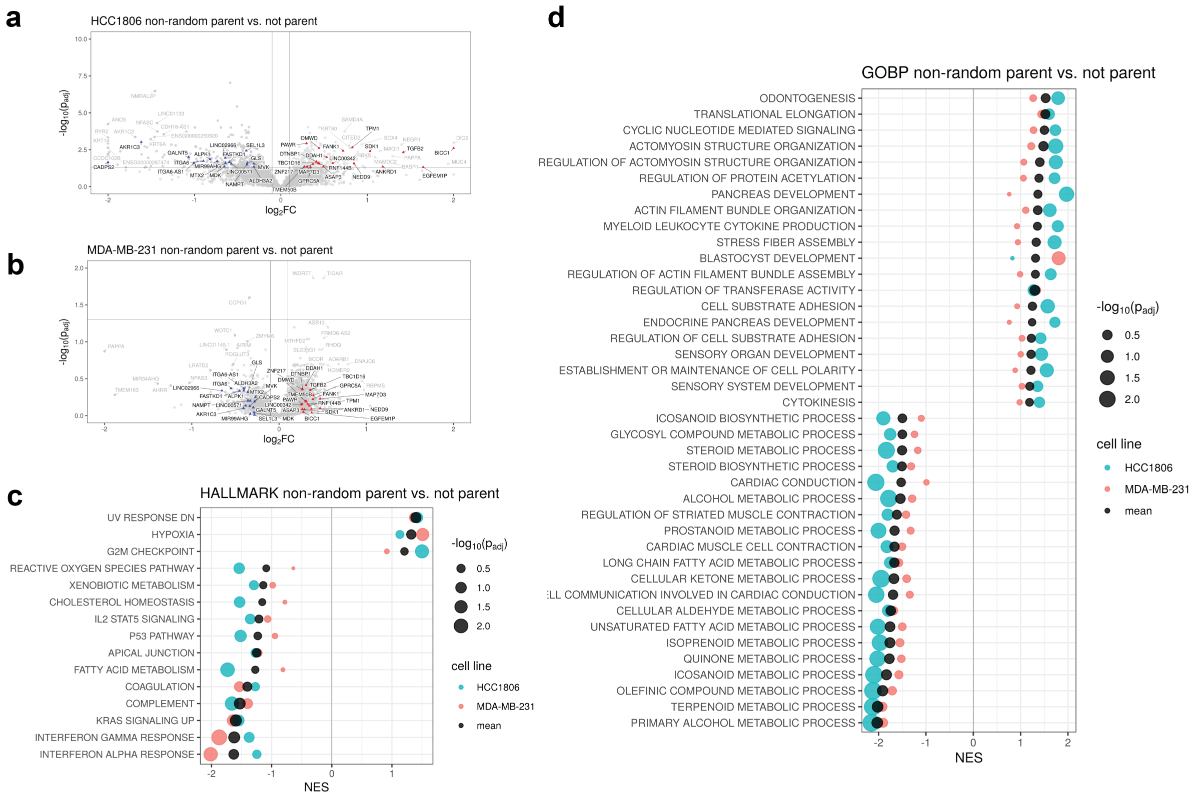


Extended Data Figure 3. Transcriptomic analysis of parental clones which occurred in fusion samples at a frequency greater than random chance. (a-b) Volcano plots showing differentially expressed genes in HCC1806, highlighting genes that are not significant in MDA-MB-231 cells but shared at log2FC > 0.2. (c) Hallmark pathways enriched in both cell lines. (d) GO Biological Process pathways enriched in both cell lines.

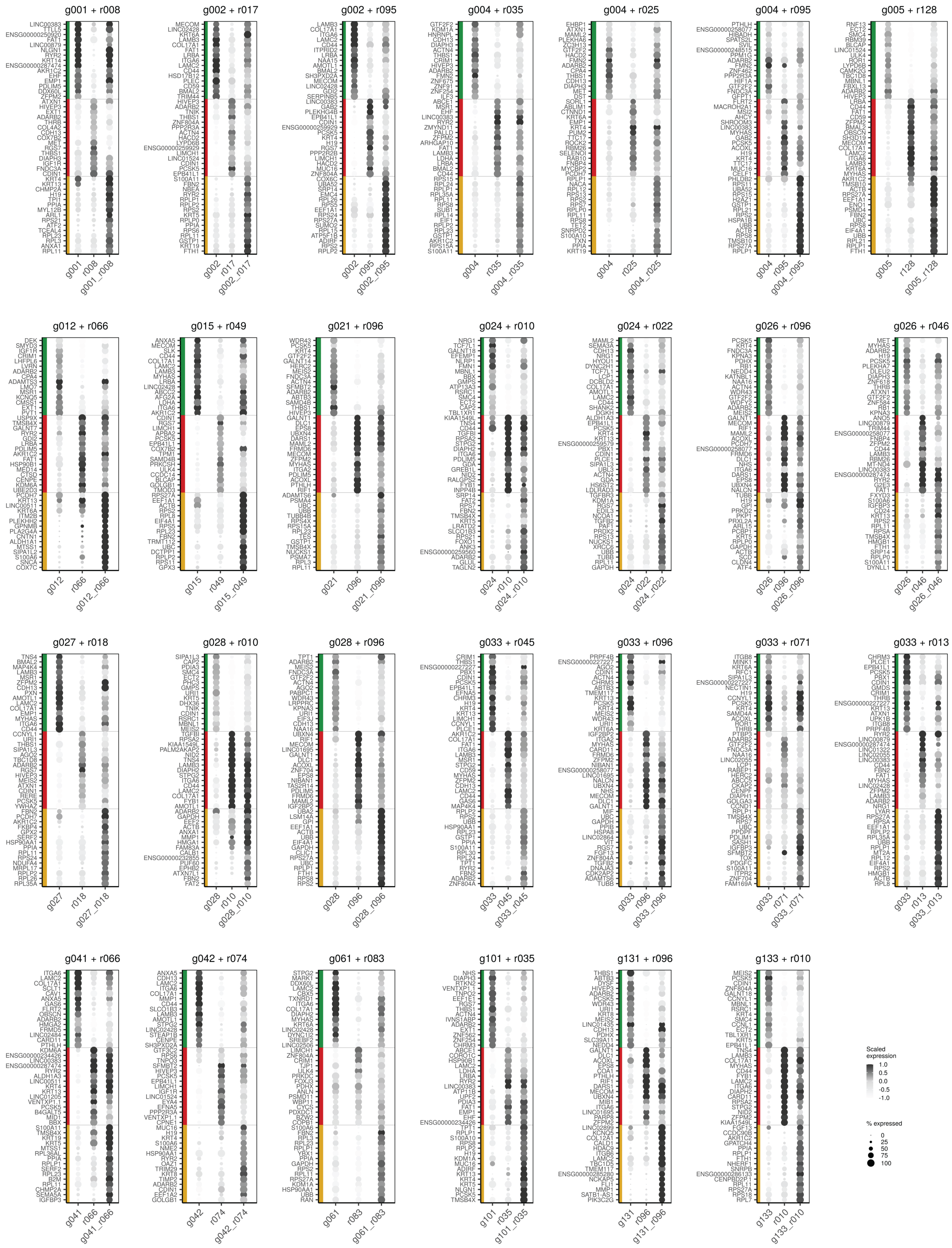

Extended Data Figure 4. Average expression of parental clone marker genes in each clone for each parent-parent-progeny set with at least 20 cells per clonal fraction in the HCC1806 population.

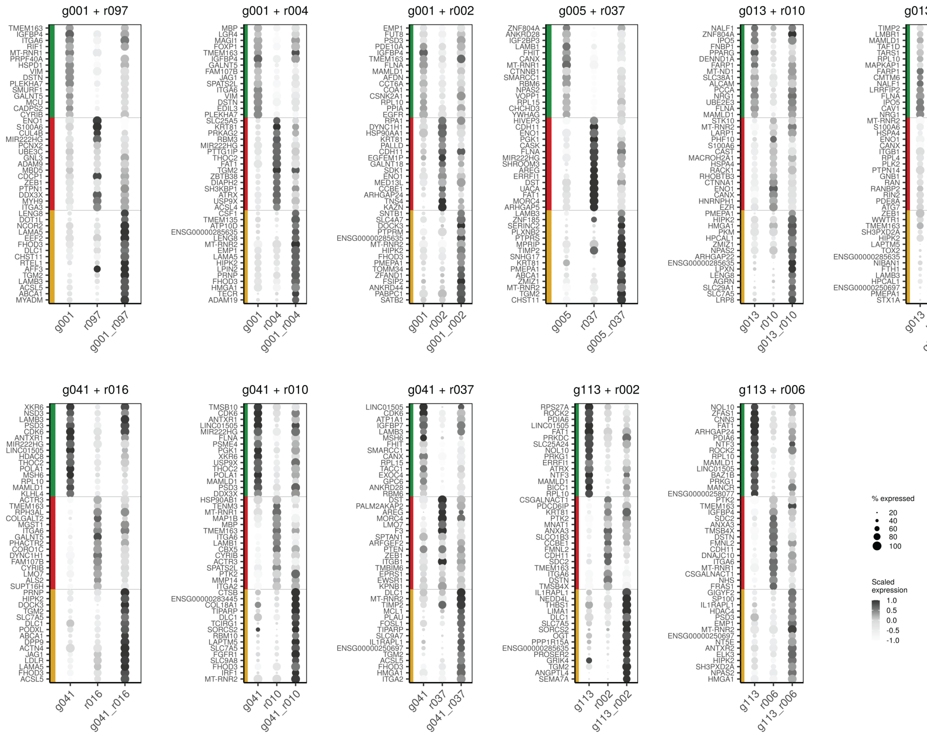


Extended Data Figure 5. Average expression of parental clone marker genes in each clone for each parent-parent-progeny set with at least 20 cells per clonal fraction in the MDA-MB-231 population.

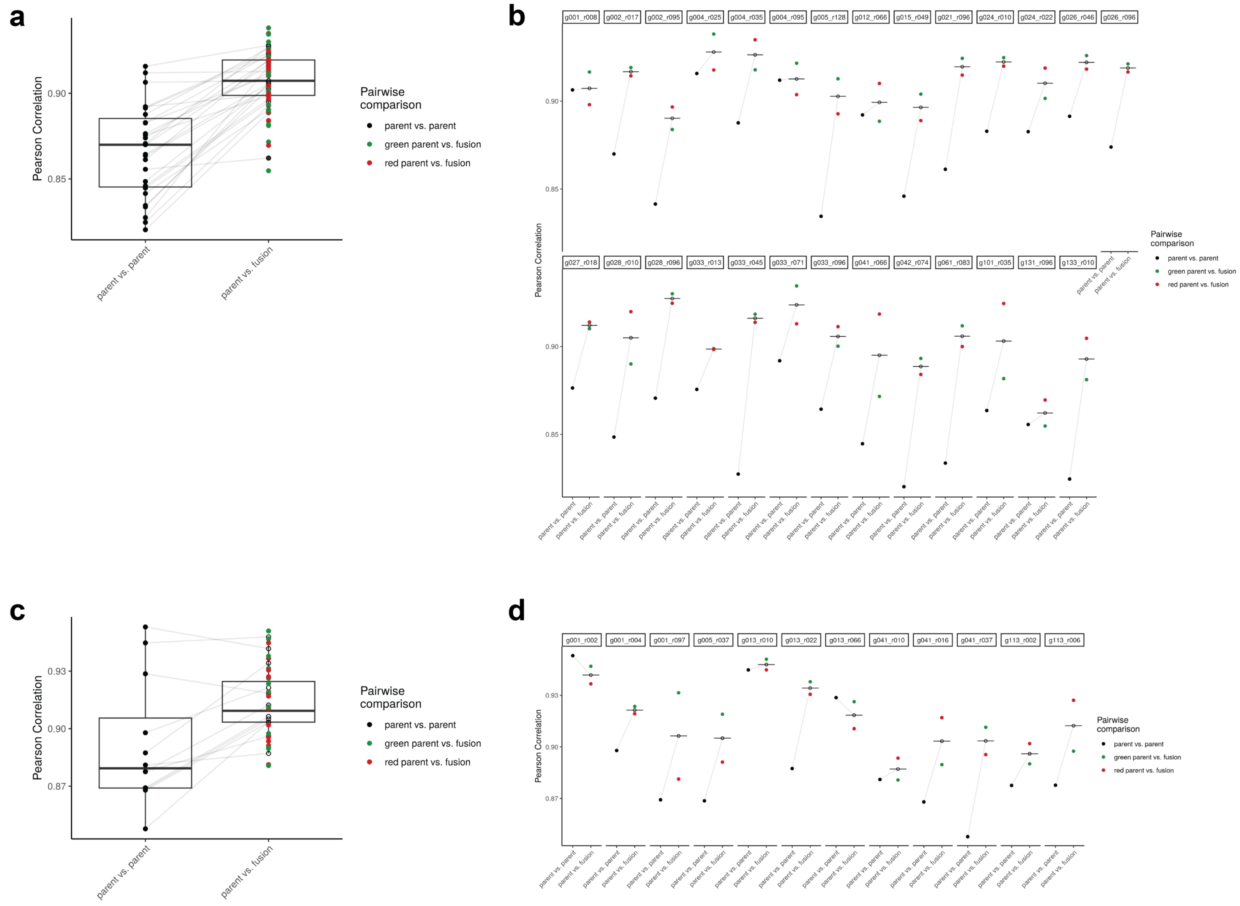


Extended Data Figure 6. Quantification of the similarity of parent-parent and parent-fusion pairs using the Pearson correlation coefficient for each parent-parent-fusion set containing a minimum of 20 cells per clonal fraction. (a) Boxplot showing the Pearson correlation coefficient for parent-parent-fusion sets. (b) Pearson correlation coefficient for each parent-parent and parent-fusion (green parent vs. fusion, and red parent vs. fusion) in each set of the HCC1806 population. (c) Boxplot summary and (d) Each parent-parent-progeny set for the MDA-MB-231 population.

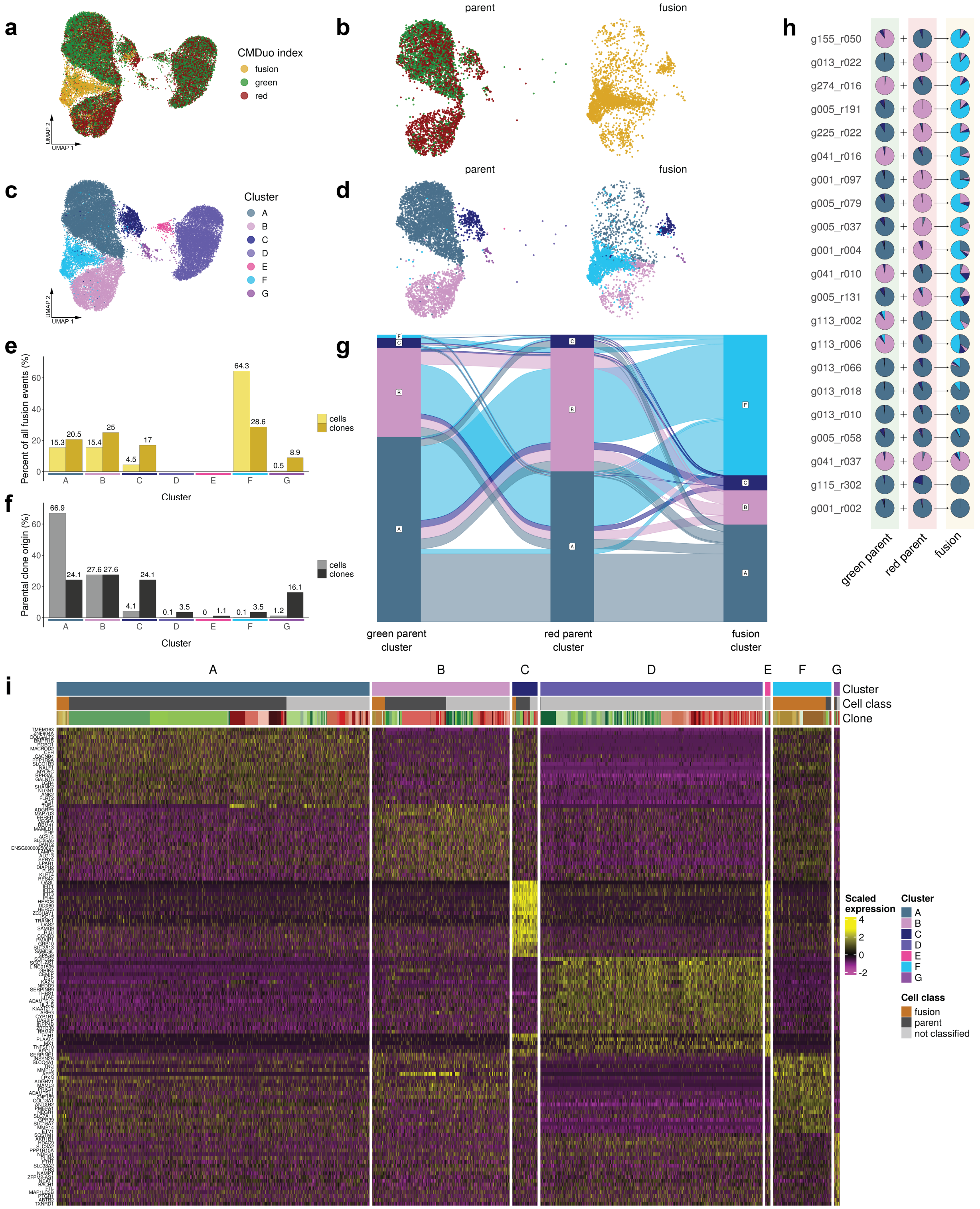


Extended Data Figure 7. There is a distinct transcriptomic state drive by cell-cell fusion. UMAP representations of all barcoded cells colored by CMDuo index (a-b) or (c-d) transcriptomic cluster in the MDA -MB-231 population. (e) Percent of all fusion events in each transcriptomic cluster separated by (left) percentage of all cells classified as fusion and (right) unique fusion clones detected. (f) Percent of all cells classified as fusion parents (left) and percent of all unique parental clones (right) found in each transcriptomic cluster. (g) Alluvial plot summarizing the relationships between parental transcriptomic clusters and fusion progeny transcriptomic cluster. (h) Pie chart representation of the proportion of each clone in each cluster for each set of parent-parent-fusion clones. (i) Heatmap showing marker gene expression for each cluster.

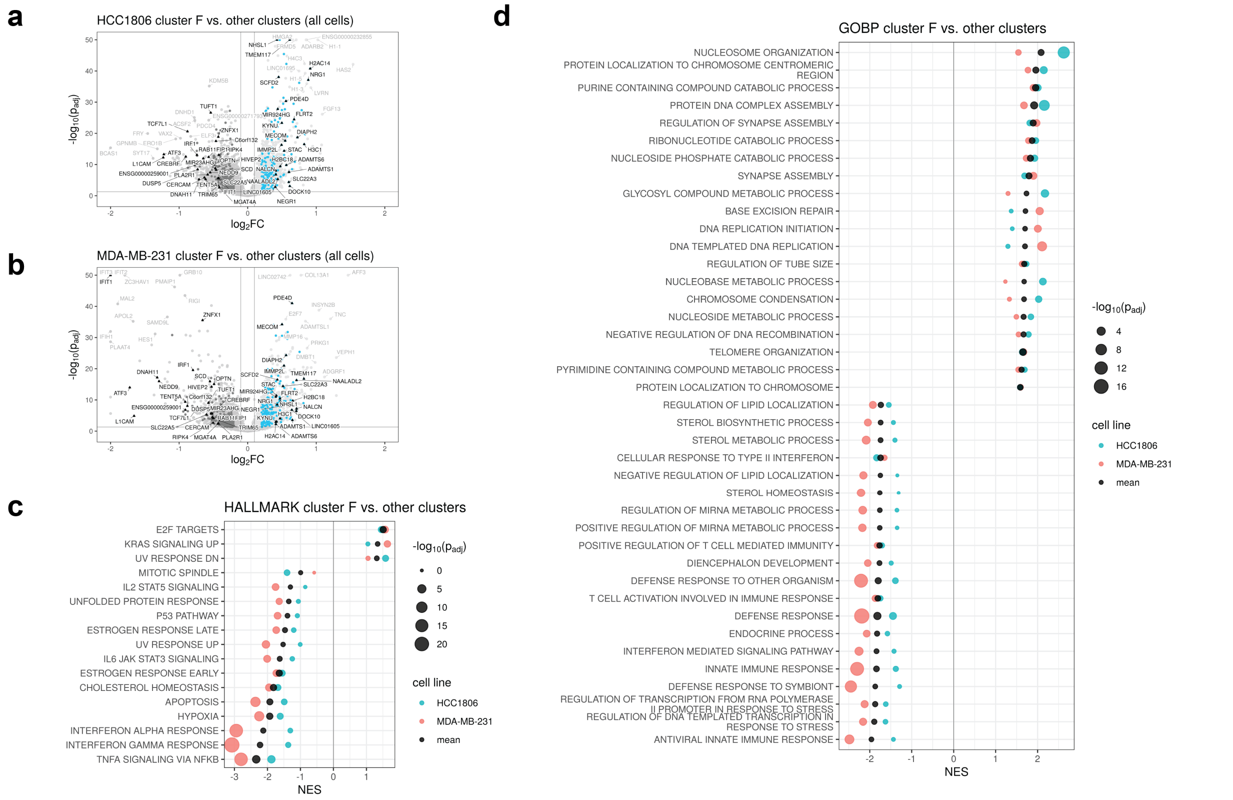


Extended Data Figure 8. Differential expression and gene set enrichment analysis of all cells in cluster F versus all cells in other clusters (a) Volcano plot of differentially expressed genes in HCC1806 and (b) MDA-MB-231 cells, highlighting the top genes differentially expressed in both cell lines. (c) Hallmark enrichment analysis showing pathways up or down in both cell lines. (d) GO biological process enrichment analysis showing pathways up or down in both cell lines.

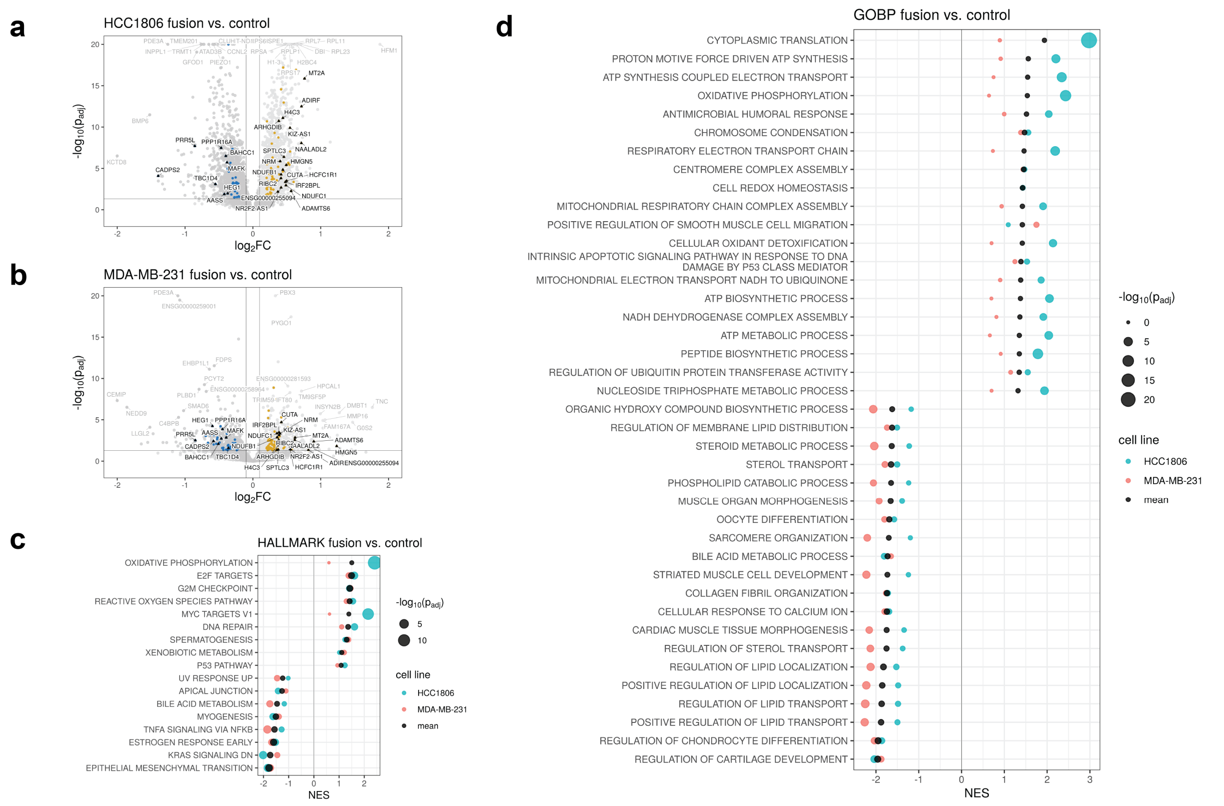


Extended Data Figure 9. Differential expression and gene set enrichment analysis of all fusion versus all control cells. (a) Volcano plot of differentially expressed genes in HCC1806 and (b) MDA-MB-231 cells, highlighting the top genes differentially expressed in both cell lines. (c) Hallmark enrichment analysis showing pathways up or down in both cell lines. (d) GO biological process enrichment analysis showing pathways up or down in both cell lines

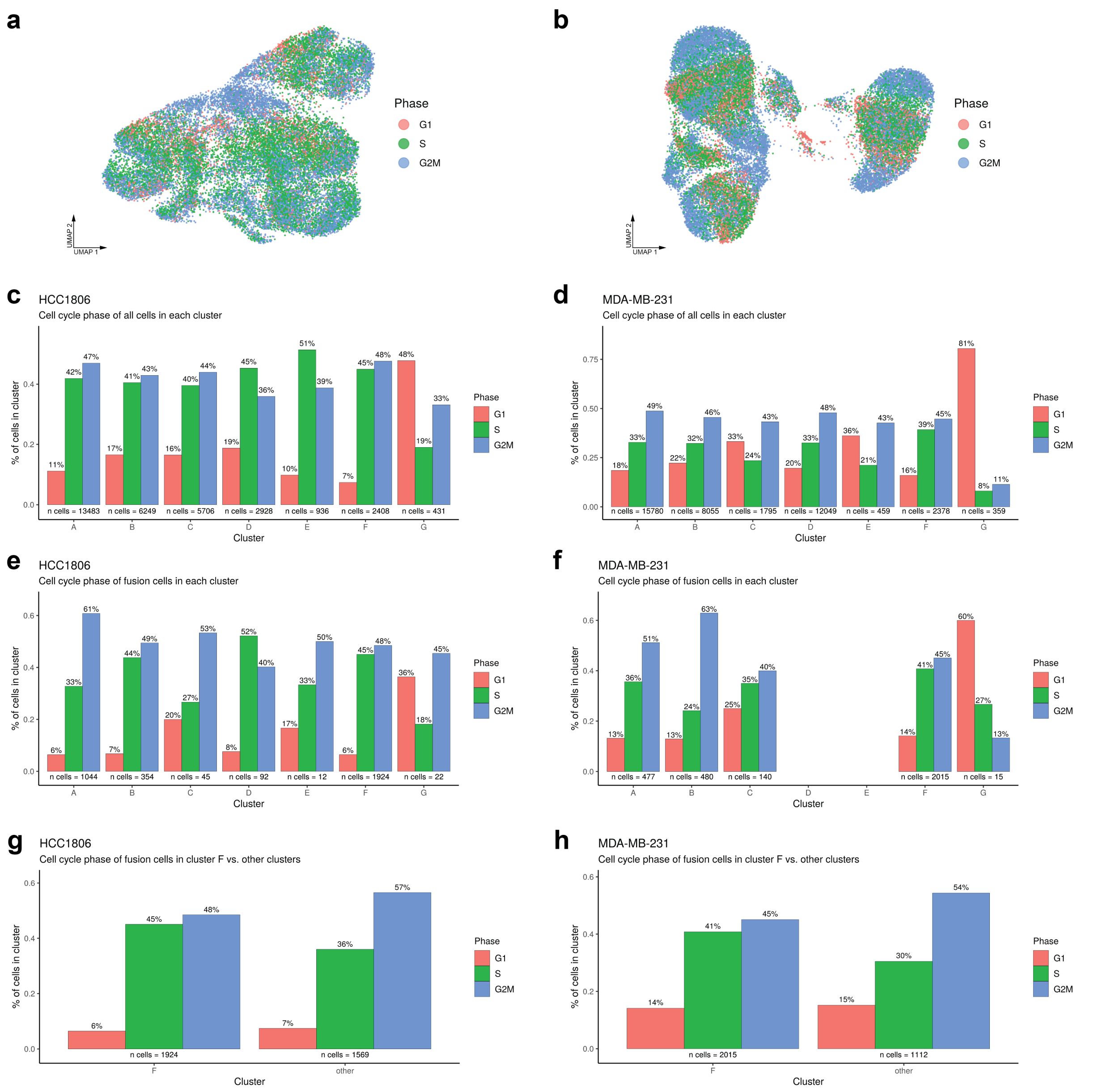


Extended Data Figure 10. Cell cycle phase patterns in each cluster. UMAP representations of the (a) HCC1806 and (b) MDA-MB-231 populations colored by cell cycle phase. Percent of all cells in each cluster in each cell cycle phase for (c) HCC1806 and (d) MDA-MB-231. Percent of fusion cells in each cluster in each cell cycle phase for (e) HCC1806 and (f) MDA-MB-231. Comparing only fusion cells in cluster F for fusion cells in other clusters for (g) HCC1806 and (h) MDA-MB-231 populations.
