## Supplemental Figures for "Mapping cell-cell fusion at single-cell resolution"

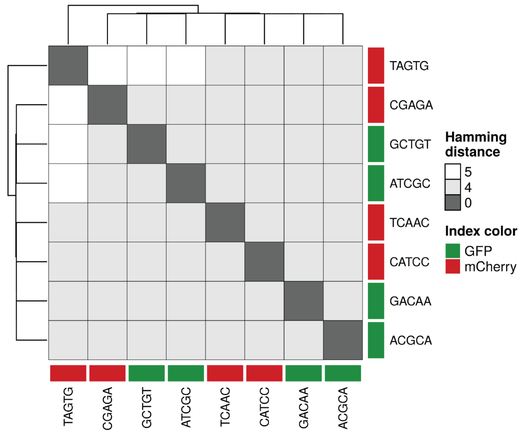


Supplemental Figure 1. Four unique 5-nt index sequences were designed for each fluorescent protein by designing sequences which maximized Hamming distance between all pairs.

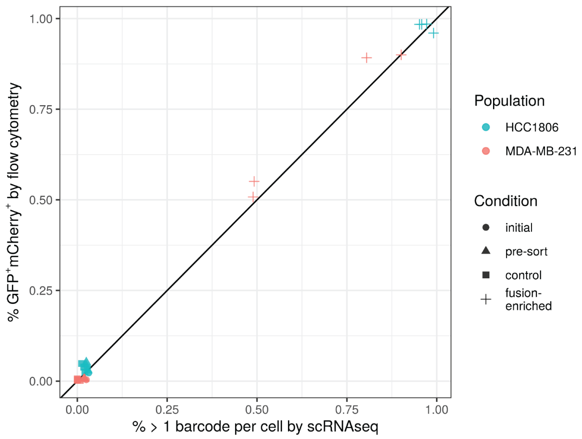


Supplemental Figure 2. Comparison of percent fusion cells in each sample with flow cytometry (% GFP+mCherry+) or CMDuo barcode assignment (% cells assigned ≥ 2 barcodes) in scRNA-seq.

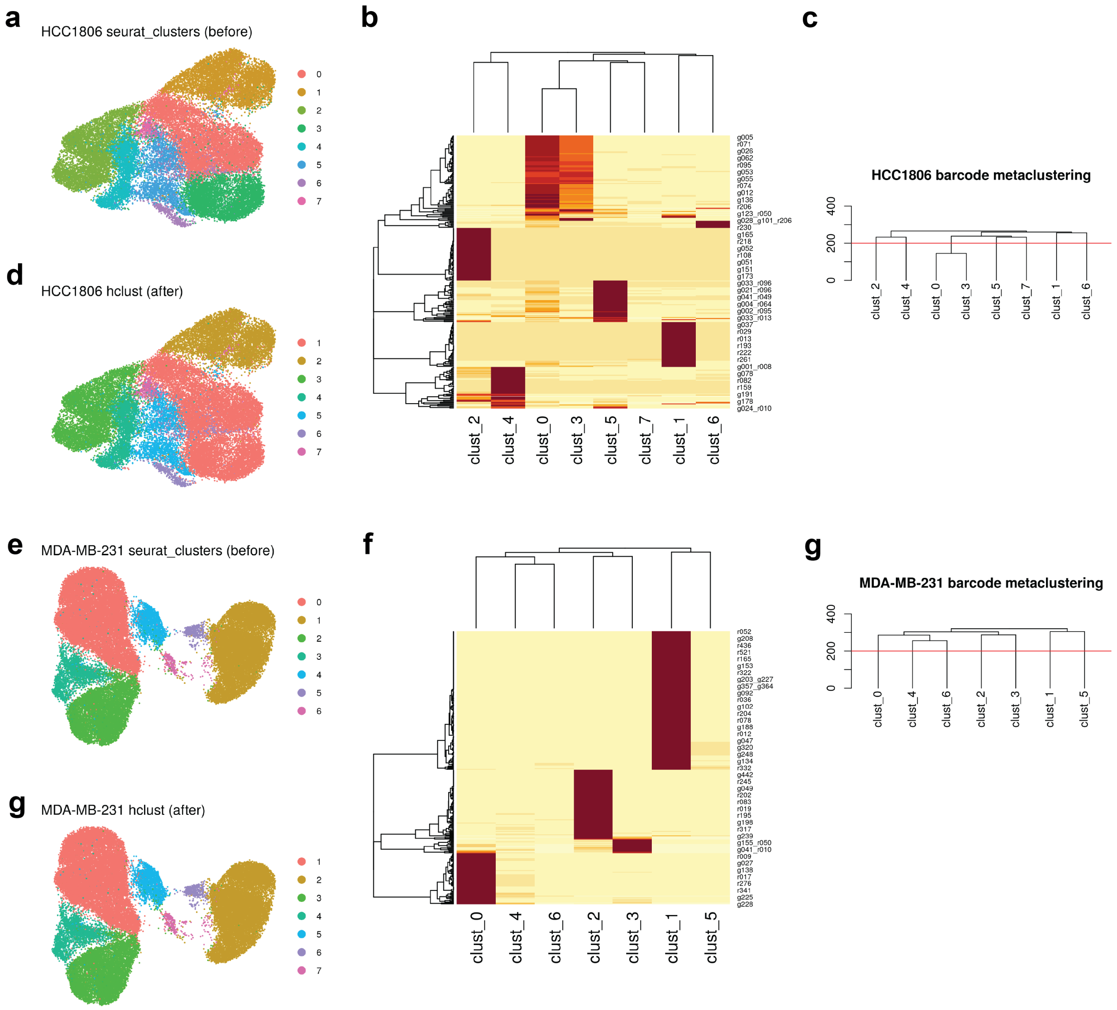


Supplemental Figure 3. Barcode-informed clustering correction. For HCC1806 (a) UMAP projection colored by Louvain clustering. (b) Heatmap showing abundance of barcodes within each cluster. (c) Distance was determined using the Canberra metric and the tree was cut at a height of 200. (d) UMAP projection after collapsing clusters 0 and 3 into one cluster. The same analysis was done for MDA-MB-231 in panels (e-g) showing no correction to the initial clustering.

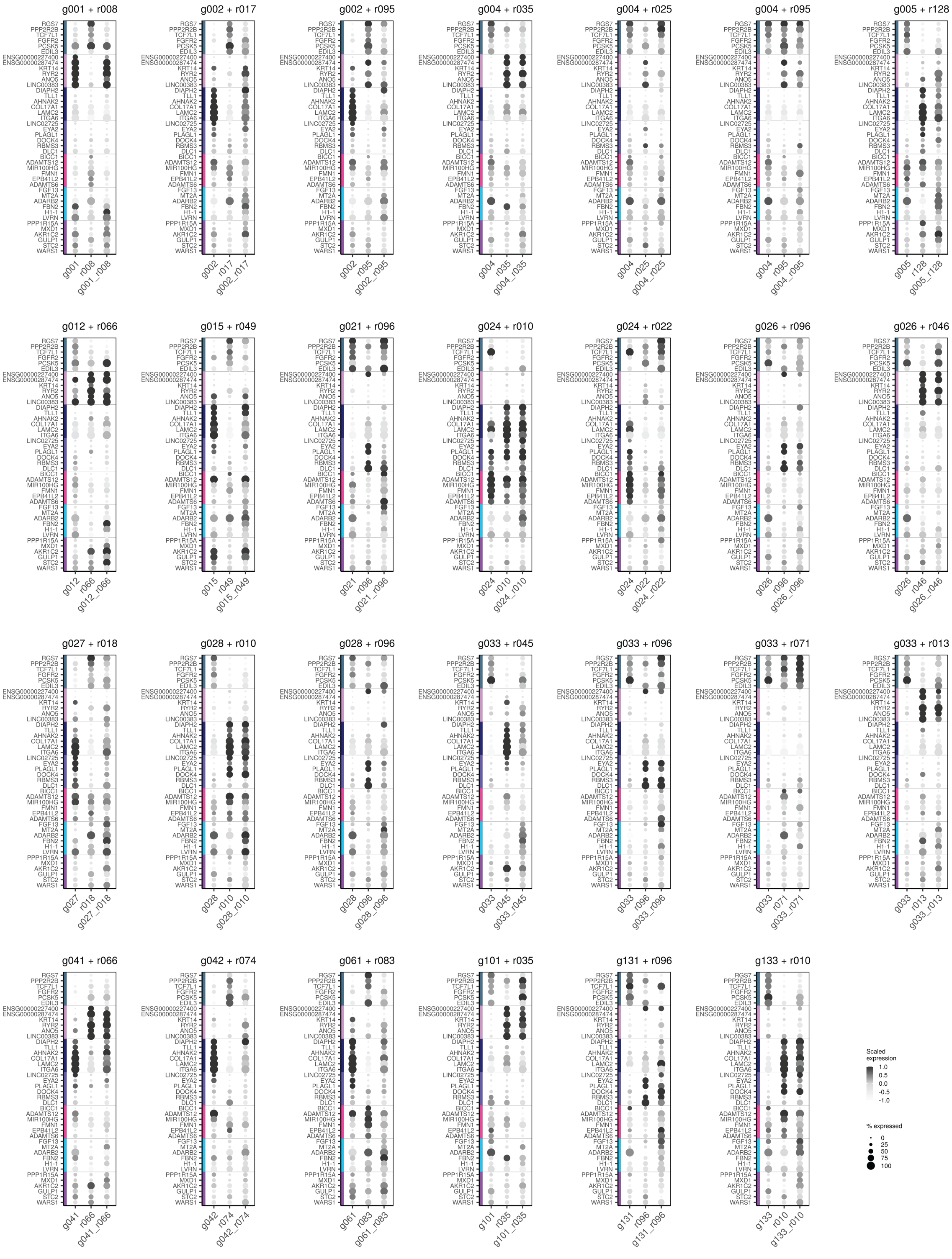


Supplemental Figure 4. Average expression of cluster marker genes in each clone for each parent-parent-progeny set with at least 20 cells per clonal fraction in the HCC1806 population.

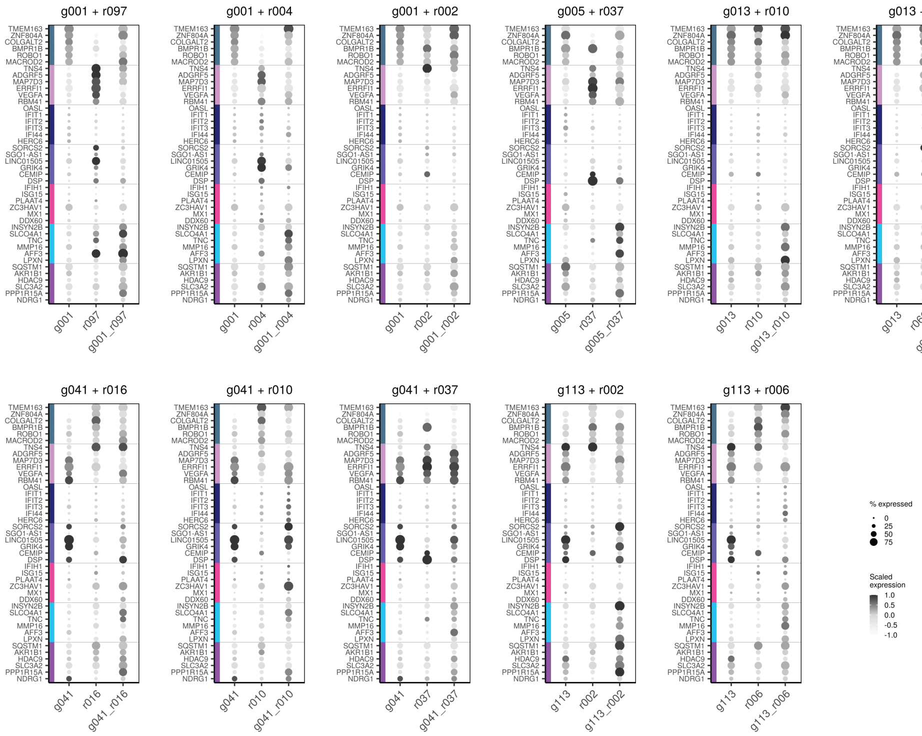


Supplemental Figure 5. Average expression of cluster marker genes in each clone for each parent-parent-progeny set with at least 20 cells per clonal fraction in the MDA-MB-231 population.

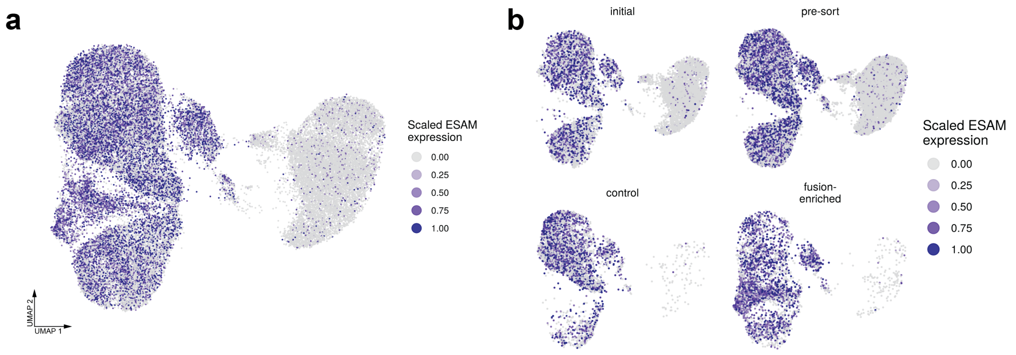


Supplemental Figure 6. ESAM expression in MDA-MB-231 cells. (a) Clusters A, B, C, and F have higher average expression of ESAM. (b) Repeated cell sorting of the MDA-MB-231 cells led to selection of the ESAM-high cells in both the control and fusion-enriched samples.
