## Supplemental Tables for "Mapping cell-cell fusion at single-cell resolution"

| | |
|---|---|
| GFP-indexed barcode 1 | GAGCCTCGTCTCCCACCGGACAANNNNNNNNNNNNNNNGTTTTGAGACGCATGCTGCA |
| GFP-indexed barcode 2 | GAGCCTCGTCTCCCACCGGCTGTNNNNNNNNNNNNNNNGTTTTGAGACGCATGCTGCA |
| GFP-indexed barcode 3 | GAGCCTCGTCTCCCACCGATCGCNNNNNNNNNNNNNNNGTTTTGAGACGCATGCTGCA |
| GFP-indexed barcode 4 | GAGCCTCGTCTCCCACCGACGCANNNNNNNNNNNNNNNGTTTTGAGACGCATGCTGCA |
| mCherry-indexed barcode 1 | GAGCCTCGTCTCCCACCGCATCCNNNNNNNNNNNNNNNGTTTTGAGACGCATGCTGCA |
| mCherry-indexed barcode 2 | GAGCCTCGTCTCCCACCGCGAGANNNNNNNNNNNNNNNGTTTTGAGACGCATGCTGCA |
| mCherry-indexed barcode 3 | GAGCCTCGTCTCCCACCGTAGTGNNNNNNNNNNNNNNNGTTTTGAGACGCATGCTGCA |
| mCherry-indexed barcode 4 | GAGCCTCGTCTCCCACCGTCAACNNNNNNNNNNNNNNNGTTTTGAGACGCATGCTGCA |

Supplemental Table 1. Indexed barcode sequences

| Cell Line | Fusion Replicate | 1st sort | | 2nd sort | | 3rd sort | | 4th sort | | final |
|---|---|---|---|---|---|---|---|---|---|---|
| | | % fusion | n sorted | % fusion | n sorted | % fusion | n sorted | % fusion | n sorted | % fusion |
| HCC1806 | 1 | 0.79 | 15,561 | 1.46 | 5,804 | 74.00 | 200,071 | - | - | 98.50 |
| HCC1806 | 2 | 1.13 | 18,207 | 1.52 | 8,605 | 76.00 | 300,098 | - | - | 98.40 |
| HCC1806 | 3 | 0.97 | 15,075 | 1.09 | 5,240 | 61.60 | 154,243 | - | - | 98.40 |
| HCC1806 | 4 | 0.75 | 18,858 | *not recorded* | 4,730 | 74.10 | 102,032 | - | - | 96.00 |
| MDA-MB-231 | 1 | 0.71 | 41,313 | 2.10 | 16,837 | 2.70 | 35,129 | 4.71 | 37,479 | 50.80 |
| MDA-MB-231 | 2 | 0.45 | 38,015 | 2.32 | 19,381 | 3.41 | 160,598 | 25.90 | 160,925 | 90.00 |
| MDA-MB-231 | 3 | 0.48 | 40,071 | 1.77 | 14,002 | 2.30 | 47,113 | 6.46 | 97,327 | 55.10 |
| MDA-MB-231 | 4 | 0.50 | 36,099 | 2.48 | 18,239 | 11.00 | 230,683 | 46.20 | 314,540 | 89.20 |

Supplemental Table 2. FACS sorting results for each fusion replicate showing % fusion at the time of sort and cell yield.

| Cell Line | condition | replicate | n_cells | n_clones | Shannon Index | Evenness |
|---|---|---|---|---|---|---|
| HCC1806 | initial | 0 | 9490 | 414 | 5.41 | 0.90 |
| HCC1806 | presort | 1 | 5015 | 259 | 4.95 | 0.89 |
| HCC1806 | presort | 2 | 6100 | 274 | 4.92 | 0.88 |
| HCC1806 | presort | 3 | 4826 | 239 | 4.84 | 0.88 |
| HCC1806 | presort | 4 | 5048 | 251 | 4.89 | 0.88 |
| HCC1806 | ctrl | 1 | 1231 | 21 | 1.53 | 0.50 |
| HCC1806 | ctrl | 2 | 1387 | 18 | 1.55 | 0.54 |
| HCC1806 | ctrl | 3 | 1506 | 24 | 1.55 | 0.49 |
| HCC1806 | ctrl | 4 | 1121 | 22 | 1.69 | 0.55 |
| HCC1806 | fusion | 1 | 1094 | 25 | 2.56 | 0.80 |
| HCC1806 | fusion | 2 | 1461 | 19 | 1.35 | 0.46 |
| HCC1806 | fusion | 3 | 1138 | 30 | 2.44 | 0.72 |
| HCC1806 | fusion | 4 | 1458 | 28 | 2.11 | 0.63 |
| MDA-MB-231 | initial | 0 | 9490 | 414 | 5.41 | 0.90 |
| MDA-MB-231 | presort | 1 | 5015 | 259 | 4.95 | 0.89 |
| MDA-MB-231 | presort | 2 | 6100 | 274 | 4.92 | 0.88 |
| MDA-MB-231 | presort | 3 | 4826 | 239 | 4.84 | 0.88 |
| MDA-MB-231 | presort | 4 | 5048 | 251 | 4.89 | 0.88 |
| MDA-MB-231 | ctrl | 1 | 1231 | 21 | 1.53 | 0.50 |
| MDA-MB-231 | ctrl | 2 | 1387 | 18 | 1.55 | 0.54 |
| MDA-MB-231 | ctrl | 3 | 1506 | 24 | 1.55 | 0.49 |
| MDA-MB-231 | ctrl | 4 | 1121 | 22 | 1.69 | 0.55 |
| MDA-MB-231 | fusion | 1 | 1094 | 25 | 2.56 | 0.80 |
| MDA-MB-231 | fusion | 2 | 1461 | 19 | 1.35 | 0.46 |
| MDA-MB-231 | fusion | 3 | 1138 | 30 | 2.44 | 0.72 |
| MDA-MB-231 | fusion | 4 | 1458 | 28 | 2.11 | 0.63 |

Supplemental Table 3. Total cell count, number of unique CMDuo clones, Shannon diversity index, and population evenness for each sample.

| Cell line | Condition | Replicate | Flow Cytometry (% $GFP^+mCherry^+$) | scRNA-seq (% ≥ 2 barcodes) |
|---|---|---|---|---|
| HCC1806 | initial | 0 | 2.28 | 3.2 |
| HCC1806 | presort | 1 | 0.83 | 2 |
| HCC1806 | presort | 2 | 5.28 | 2.5 |
| HCC1806 | presort | 3 | 4.46 | 2.4 |
| HCC1806 | presort | 4 | 2.89 | 2.9 |
| HCC1806 | ctrl | 1 | 4.85 | 1.2 |
| HCC1806 | ctrl | 2 | 3.70 | 2.8 |
| HCC1806 | ctrl | 3 | 3.86 | 1.8 |
| HCC1806 | ctrl | 4 | 2.50 | 2.1 |
| HCC1806 | fusion | 1 | 98.50 | 97.2 |
| HCC1806 | fusion | 2 | 98.40 | 95.2 |
| HCC1806 | fusion | 3 | 98.40 | 95.8 |
| HCC1806 | fusion | 4 | 96.00 | 99.1 |
| MDA-MB-231 | initial | 0 | 0.39 | 2.5 |
| MDA-MB-231 | presort | 1 | 0.54 | 2.1 |
| MDA-MB-231 | presort | 2 | 0.52 | 1.8 |
| MDA-MB-231 | presort | 3 | 0.71 | 2 |
| MDA-MB-231 | presort | 4 | 0.67 | 1.8 |
| MDA-MB-231 | ctrl | 1 | 0.41 | 0 |
| MDA-MB-231 | ctrl | 2 | 0.22 | 0 |
| MDA-MB-231 | ctrl | 3 | 0.22 | 0.8 |
| MDA-MB-231 | ctrl | 4 | 0.58 | 0 |
| MDA-MB-231 | fusion | 1 | 50.80 | 48.9 |
| MDA-MB-231 | fusion | 2 | 90.00 | 90.1 |
| MDA-MB-231 | fusion | 3 | 55.10 | 49.2 |
| MDA-MB-231 | fusion | 4 | 89.20 | 80.5 |

Supplemental Table 4. Quantification of percent fusion cells in each sample with flow cytometry (% GFP+mCherry+) or CMDuo barcode assignment (% cells assigned ≥ 2 barcodes) in scRNA-seq.

| | | |
|---|---|---|
| Highly selected parent clones | Up | BICC1, EGFEM1P, TGFB2, ANKRD1, SDK1, NEDD9, TPM1, SAMD4A, FANK1, LINC00342, RNF213-AS1, MYLIP, CRIM1, DDAH1, UGCG, ASAP3, CAP2, ZNF347, TMEM156, RNF144B, DMWD, TBC1D16, DTNBP1, CCN1, ZNF217, TCF7L2, ANO6, STRBP, GPRC5A, SP110, ALMS1, CNPY3, MAP7D3, PAWR, TMEM50B, R3HCC1L, LITAF |
| | Down | CADPS2, AKR1C3, GALNT5, ITGA6-AS1, ITGA6, ALPK1, MIR99AHG, MDK, LINC02966, MTX2, FASTKD1, LINC00571, SEL1L3, NAMPT, GLS, MVK, ALDH3A2 |

Supplemental Table 5. Differentially expressed genes in highly selected parental clones (abs(log2FC) > 0.2 and p-adj <=0.05 in HCC1806; abs(log2FC) > 0.2 and p-adj <=1 in MDA-MB-231)

| | |
|---|---|
| A | RGS7, PPP2R2B, TCF7L1, FGFR2, PCSK5, EDIL3, FRAS1, EEF1A2, PLEKHG4B, CCNYL1, MEGF6, THBS1, ERRFI1, MEST, CLDN11, ANK1, CRIM1, H19, HIVEP3, LINC01224, WNT7B, TPM1, PLD1, SAMD12, CDIN1, LYPD6B, LIMCH1, ADARB2, ZNF804A, CHRM3 |
| B | ENSG00000227400, ENSG00000287474, KRT14, RYR2, ANO5, LINC00383, LINC00879, KRT13, KRT4, NLGN1, CAMK1D, ENSG00000250920, FBN2, MTSS1, MUC16, PKP1, TET2, LINC01322, EHF, LINC00511, NFKBIZ, H19, AKR1C2, IGFBP3, ENSG00000234426, KLF5, ARHGAP26, LINC01205, FOXN3, MYEOV |
| C | DIAPH2, TLL1, AHNAK2, COL17A1, LAMC2, ITGA6, HDAC9, STPG2, SLC7A11, LAMA3, CDH13, ANXA5, LINC01322, DDX60L, KRT6A, TGFBI, LAMB3, LINC02484, EGFP, BLTP1, SCLT1, SLCO1B3, ENSG00000286797, NET1, INPP4B, RPSA2, CALD1, NID2, AKR1C2, SCD |
| D | LINC02725, EYA2, PLAGL1, DOCK4, RBMS3, DLC1, DAPK1, ADAMTS12, LAMA4, KIAA1549L, ENSG00000289349, ITGA6, LINC00879, GALNT1, DTNB, VIT, STPG2, AHNAK2, LAMC2, BCAR3, MECOM, LRP1B, RPSA2, ENSG00000248515, TNS4, CAPRIN2, MAP4K4, TIMP3, LINC01695, IMMP2L |
| E | BICC1, ADAMTS12, TCF7L1, MIR100HG, FMN1, EPB41L2, ADAMTS6, ENSG00000229855, P3H2, COL27A1, CLDN1, PRSS23, MICAL2, PALM2AKAP2, TPM1, NRG1, TGFB2, PHLDB2, PDZD2, SHANK2, JAZF1, NR2F1-AS1, LCP1, NHS, CDH13, FHOD3, NALCN, SLC1A3, SEMA3A, PAM |
| F | FGF13, MT2A, DIAPH2, ADARB2, FBN2, H1-1, LVRN, S100A2, H4C3, DKK1, H2BC4, H1-5, KRT6A, SEMA3D, FUT9, MIF, H1-3, UBB, RPS15, UBA52, HSPE1, H1-4, KDM1A, ENSG00000232855, PSMB6, FAU, ADAMTS6, RPS21, RPS27A, GSTP1 |
| G | PPP1R15A, MXD1, AKR1C2, GULP1, STC2, WARS1, ENSG00000291336, NEAT1, SQSTM1, NDRG1, GFPT1, LINC00662, CD55, SEL1L, PCF11, SLC7A11, PPP1R15B, SARS1, NAMPT, OPTN, IFI16, GCLC, CCDC186, MALAT1, SPTY2D1, GARS1, ABTB2, NR1D2, RB1CC1, RAB5A |

Supplemental Table 6. HCC1806 cluster marker genes

| | |
|---|---|
| A | TMEM163, ZNF804A, COLGALT2, BMPR1B, ROBO1, MACROD2, CFH, CACNB4, PPP1R9A, SLCO1B3, NALF1, MYO5C, RPH3AL, GALNT5, LGR4, SHANK2, NLGN1, ANK2, FLRT2, JAG1, PPARG, IGFBP4, CSGALNACT1, FAM107B, CADPS2, PEG10, MGST1, PXDN, FOXP1, ANKRD44 |
| B | TNS4, ADGRF5, MAP7D3, ERRFI1, VEGFA, RBM41, MAMLD1, EHF, ACSL4, SLC25A5, DANT2, ENSG00000250697, LAMP2, ALG13, SPRY4, LPAR1, DIAPH2, PLS3, KLHL4, RPS4X, MORF4L2, PLAU, THOC2, SDK1, ELL2, S100A6, LINC00511, PGK1, NRP1, STAG2 |
| C | OASL, IFIT1, IFIT2, IFIT3, IFI44, HERC6, DDX60, HERC5, ZC3HAV1, ISG15, TRANK1, OAS2, SAMD9, RIGI, CCND3, PMAIP1, GRB10, SLC2A13, SAMD9L, SPAG9, TTC28, HIP1R, B4GALT5, VEGFC, FNDC3A, PARP14, CHN2, JAK2, MAPRE2, ZNFX1 |
| D | SORCS2, SGO1-AS1, LINC01505, GRIK4, CEMIP, DSP, KAZN, NEDD9, SERPINB9, THBS1, LITAF, ADAMTS12, HLA-B, KIAA1217, AREG, CYP1B1, DAB2IP, INPP4B, ZBTB38, RBM47, CD82, EPAS1, TTC39C, BACE2, MED14, TLE1, GLIS3, HLA-C, WWC1, LYST |
| E | IFIH1, IFIT3, IFIT2, IFIT1, ISG15, OASL, PLAAT4, ZC3HAV1, MX1, HERC6, DDX60, HERC5, IFI44, RIGI, SAMD9L, TRANK1, SAMD9, TNFSF10, APOL1, SERPINE1, WARS1, PARP14, PMAIP1, HIP1R, SPAG9, DDX60L, CCND3, DUSP10, B4GALT5, GRB10 |
| F | INSYN2B, SLCO4A1, TNC, MMP16, AFF3, LPXN, ADGRV1, MAML3, PRKG1, ADAMTSL1, ZNF185, COL13A1, ANTXR2, PMEPA1, NEGR1, SLC7A11, GPR39, SLC16A7, MMP14, ETV1, PITPNC1, NALF1, NAALADL2, ANKRD44, INPP5A, SEMA3A, ENSG00000285635, DOCK3, GPAT3, GINS4 |
| G | SQSTM1, AKR1B1, HDAC9, SLC3A2, PPP1R15A, NDRG1, PLIN2, SLC7A11, FTH1, SLC38A2, IER3, NAMPT, ZFPM2-AS1, NEAT1, BACH1, FTL, MAP1LC3B, PTGR1, ABTB2, TXNRD1, CPEB2, ERRFI1, SLC38A1, UGDH, RB1CC1, ATP6V1C1, HIVEP2, GBE1, SPIRE1, HIF1A-AS3 |

Supplemental Table 7. MDA-MB-231 cluster marker genes

| gene | HCC1 806 | MDA-MB-231 | mean_log2FC | | gene | HCC1 806 | MDA-MB-231 | mean_log2FC | | gene | HCC1 806 | MDA-MB-231 | mean_log2FC |
|---|---|---|---|---|---|---|---|---|---|---|---|---|---|
| H2AC14 | 0.912 | 0.41 | 0.912 | | CRPPA | 0.375 | 0.534 | 0.375 | | ADGRA3 | 0.264 | 0.283 | 0.264 |
| NRG1 | 0.884 | 0.43 | 0.884 | | COMMD10 | 0.371 | 0.248 | 0.371 | | RRM1 | 0.263 | 0.212 | 0.263 |
| HTR7 | 0.837 | 0.637 | 0.837 | | SERPINE2 | 0.371 | 0.567 | 0.371 | | ADK | 0.262 | 0.291 | 0.262 |
| H4C3 | 0.829 | 0.357 | 0.829 | | ENSG00000 287621 | 0.365 | 0.425 | 0.365 | | MZF1 | 0.261 | 0.276 | 0.261 |
| H3C1 | 0.824 | 0.473 | 0.824 | | GPHN | 0.361 | 0.318 | 0.361 | | CEP162 | 0.258 | 0.224 | 0.258 |
| H2AC20 | 0.787 | 0.294 | 0.787 | | GALNT10 | 0.361 | 0.282 | 0.361 | | DNAJC19 | 0.258 | 0.331 | 0.258 |
| H1-4 | 0.752 | 0.283 | 0.752 | | PIP4P2 | 0.361 | 0.227 | 0.361 | | HDAC8 | 0.257 | 0.343 | 0.257 |
| DIAPH2 | 0.725 | 0.561 | 0.725 | | SYT1 | 0.36 | 0.581 | 0.36 | | TRMT11 | 0.256 | 0.265 | 0.256 |
| FRMD5 | 0.708 | 0.303 | 0.708 | | EYA4 | 0.357 | 0.235 | 0.357 | | SLC4A7 | 0.256 | 0.204 | 0.256 |
| H2AC11 | 0.701 | 0.626 | 0.701 | | ARHGAP18 | 0.356 | 0.345 | 0.356 | | ATXN7L1 | 0.256 | 0.26 | 0.256 |
| MT1E | 0.687 | 0.332 | 0.687 | | FAM111B | 0.353 | 0.355 | 0.353 | | LINC01376 | 0.255 | 0.344 | 0.255 |
| FLRT2 | 0.677 | 0.472 | 0.677 | | CNTNAP3P2 | 0.352 | 0.631 | 0.352 | | TRAPPC9 | 0.254 | 0.413 | 0.254 |
| H1-2 | 0.669 | 0.351 | 0.669 | | DYNC2H1 | 0.351 | 0.465 | 0.351 | | TMEM106C | 0.252 | 0.369 | 0.252 |
| KCNQ5 | 0.658 | 0.281 | 0.658 | | CDK4 | 0.351 | 0.299 | 0.351 | | PPP2R3C | 0.251 | 0.279 | 0.251 |
| HMGA2 | 0.655 | 0.289 | 0.655 | | DCBLD2 | 0.351 | 0.246 | 0.351 | | YEATS4 | 0.249 | 0.327 | 0.249 |
| NAV3 | 0.647 | 0.756 | 0.647 | | CMSS1 | 0.349 | 0.304 | 0.349 | | TMEM38B | 0.248 | 0.206 | 0.248 |
| DOCK4 | 0.639 | 0.445 | 0.639 | | PRDM5 | 0.347 | 0.281 | 0.347 | | H2BC14 | 0.247 | 0.813 | 0.247 |
| DOCK10 | 0.615 | 0.71 | 0.615 | | MT2A | 0.345 | 0.243 | 0.345 | | TBC1D5 | 0.244 | 0.354 | 0.244 |
| TMEM117 | 0.614 | 0.82 | 0.614 | | PRICKLE1 | 0.345 | 0.297 | 0.345 | | TBC1D19 | 0.243 | 0.311 | 0.243 |
| ADAMTS1 | 0.603 | 0.652 | 0.603 | | LINC00342 | 0.344 | 0.299 | 0.344 | | PRIM1 | 0.243 | 0.438 | 0.243 |
| SLC22A3 | 0.599 | 0.515 | 0.599 | | H2BC21 | 0.343 | 0.531 | 0.343 | | ZBTB20 | 0.243 | 0.284 | 0.243 |
| NR2F1-AS1 | 0.578 | 0.319 | 0.578 | | ARSB | 0.341 | 0.392 | 0.341 | | PLCL2 | 0.242 | 0.218 | 0.242 |
| HIVEP3 | 0.564 | 0.291 | 0.564 | | SLC10A7 | 0.341 | 0.232 | 0.341 | | EEF1AKMT1 | 0.24 | 0.27 | 0.24 |
| ADAMTS6 | 0.563 | 0.412 | 0.563 | | MAML2 | 0.34 | 0.265 | 0.34 | | POLR3G | 0.239 | 0.309 | 0.239 |
| PDE4D | 0.558 | 0.647 | 0.558 | | ULK4 | 0.338 | 0.263 | 0.338 | | FAM13A | 0.237 | 0.273 | 0.237 |
| MECOM | 0.547 | 0.501 | 0.547 | | H2AC17 | 0.336 | 0.608 | 0.336 | | PCNX2 | 0.236 | 0.368 | 0.236 |
| CHST11 | 0.542 | 0.283 | 0.542 | | DPY19L1 | 0.334 | 0.256 | 0.334 | | PBX1 | 0.235 | 0.243 | 0.235 |
| SLC7A11 | 0.539 | 0.508 | 0.539 | | H2BC3 | 0.331 | 0.271 | 0.331 | | SKA3 | 0.235 | 0.251 | 0.235 |
| H3C3 | 0.535 | 0.664 | 0.535 | | ENSG00000 283415 | 0.327 | 0.367 | 0.327 | | CCNE2 | 0.234 | 0.378 | 0.234 |
| STAC | 0.532 | 0.419 | 0.532 | | CAMKMT | 0.323 | 0.335 | 0.323 | | KCTD1 | 0.233 | 0.471 | 0.233 |
| ITGBL1 | 0.531 | 0.257 | 0.531 | | AGA-DT | 0.32 | 0.486 | 0.32 | | PHKA1 | 0.233 | 0.451 | 0.233 |
| LINC00941 | 0.529 | 0.212 | 0.529 | | RTKN2 | 0.32 | 0.281 | 0.32 | | MDM1 | 0.233 | 0.482 | 0.233 |
| GPR39 | 0.529 | 0.584 | 0.529 | | HMGA2-AS1 | 0.319 | 0.321 | 0.319 | | TFB1M | 0.232 | 0.233 | 0.232 |
| TNIK | 0.522 | 0.338 | 0.522 | | ENSG00000 227598 | 0.316 | 0.257 | 0.316 | | LANCL2 | 0.231 | 0.206 | 0.231 |
| INSYN2B | 0.521 | 1.039 | 0.521 | | PLEK2 | 0.314 | 0.285 | 0.314 | | MTHFD1L | 0.229 | 0.288 | 0.229 |
| CDIN1 | 0.517 | 0.252 | 0.517 | | WWOX | 0.314 | 0.434 | 0.314 | | RANBP17 | 0.227 | 0.301 | 0.227 |
| LPAR1 | 0.499 | 0.237 | 0.499 | | SDK1 | 0.313 | 0.273 | 0.313 | | DIAPH3 | 0.227 | 0.227 | 0.227 |
| KYNU | 0.499 | 0.422 | 0.499 | | FHOD3 | 0.312 | 0.294 | 0.312 | | LAMA3 | 0.226 | 0.502 | 0.226 |
| IMMP2L | 0.495 | 0.53 | 0.495 | | BMAL2 | 0.308 | 0.219 | 0.308 | | PRH1 | 0.225 | 0.465 | 0.225 |
| GPAT3 | 0.49 | 0.49 | 0.49 | | CRADD | 0.307 | 0.279 | 0.307 | | ENSG00000 225218 | 0.224 | 0.295 | 0.224 |
| CHAC2 | 0.486 | 0.322 | 0.486 | | CEP128 | 0.305 | 0.283 | 0.305 | | APEX2 | 0.223 | 0.246 | 0.223 |
| COL13A1 | 0.48 | 0.777 | 0.48 | | SAMD1 | 0.303 | 0.249 | 0.303 | | ME3 | 0.223 | 0.271 | 0.223 |
| ARHGEF4 | 0.476 | 0.204 | 0.476 | | MNS1 | 0.298 | 0.254 | 0.298 | | RMI1 | 0.221 | 0.236 | 0.221 |
| H2BC18 | 0.475 | 0.658 | 0.475 | | FARS2 | 0.297 | 0.201 | 0.297 | | SH3RF2 | 0.22 | 0.223 | 0.22 |
| FUT8 | 0.468 | 0.356 | 0.468 | | COL4A5 | 0.297 | 0.464 | 0.297 | | SVIP | 0.218 | 0.299 | 0.218 |
| PRKCA | 0.468 | 0.382 | 0.468 | | CDKAL1 | 0.296 | 0.286 | 0.296 | | PARPBP | 0.217 | 0.363 | 0.217 |
| NHS | 0.462 | 0.291 | 0.462 | | CAPRIN2 | 0.294 | 0.234 | 0.294 | | MFSD1 | 0.215 | 0.241 | 0.215 |
| SCFD2 | 0.451 | 0.433 | 0.451 | | ZNF804A | 0.292 | 0.415 | 0.292 | | GMDS | 0.214 | 0.346 | 0.214 |
| H3C12 | 0.451 | 0.221 | 0.451 | | LRR1 | 0.292 | 0.362 | 0.292 | | TIAM1 | 0.214 | 0.553 | 0.214 |
| ENSG00000 258077 | 0.444 | 0.23 | 0.444 | | VWA8 | 0.288 | 0.3 | 0.288 | | SRGAP1 | 0.213 | 0.309 | 0.213 |
| ABCC4 | 0.443 | 0.307 | 0.443 | | NSMAF | 0.286 | 0.359 | 0.286 | | BCKDHB | 0.213 | 0.233 | 0.213 |
| LINC01605 | 0.439 | 0.644 | 0.439 | | RAP1GDS1 | 0.286 | 0.203 | 0.286 | | COPS7A | 0.213 | 0.217 | 0.213 |
| MIR924HG | 0.439 | 0.411 | 0.439 | | PLCB1 | 0.286 | 0.305 | 0.286 | | RSRC1 | 0.212 | 0.255 | 0.212 |
| MEIS2 | 0.437 | 0.343 | 0.437 | | LINC01572 | 0.285 | 0.376 | 0.285 | | UBE2E3 | 0.212 | 0.227 | 0.212 |
| DYNC1I1 | 0.428 | 0.618 | 0.428 | | MSRA | 0.284 | 0.477 | 0.284 | | TRIM59-IFT80 | 0.212 | 0.369 | 0.212 |
| NBEA | 0.427 | 0.318 | 0.427 | | FBXO5 | 0.283 | 0.309 | 0.283 | | DCLRE1B | 0.21 | 0.207 | 0.21 |
| NHSL1 | 0.426 | 0.477 | 0.426 | | ENOX2 | 0.282 | 0.205 | 0.282 | | SHMT2 | 0.208 | 0.232 | 0.208 |
| TMEM14A | 0.418 | 0.209 | 0.418 | | H2BC12 | 0.281 | 0.413 | 0.281 | | ITFG1 | 0.208 | 0.331 | 0.208 |
| KLF12 | 0.415 | 0.258 | 0.415 | | H3C2 | 0.28 | 0.684 | 0.28 | | RFX3 | 0.207 | 0.326 | 0.207 |
| NAALADL2 | 0.412 | 0.712 | 0.412 | | IGFBP7 | 0.278 | 0.357 | 0.278 | | AGAP1 | 0.206 | 0.262 | 0.206 |
| NALCN | 0.411 | 0.713 | 0.411 | | ITGA2 | 0.276 | 0.215 | 0.276 | | POLQ | 0.205 | 0.498 | 0.205 |
| H2AC4 | 0.409 | 0.355 | 0.409 | | CNTNAP3 | 0.275 | 0.973 | 0.275 | | TEX9 | 0.203 | 0.291 | 0.203 |
| CNPY2 | 0.409 | 0.214 | 0.409 | | UBE3D | 0.274 | 0.346 | 0.274 | | LRBA | 0.203 | 0.221 | 0.203 |
| NEGR1 | 0.402 | 0.478 | 0.402 | | TMC7 | 0.272 | 0.319 | 0.272 | | BTBD9 | 0.201 | 0.277 | 0.201 |
| DPYD | 0.399 | 0.224 | 0.399 | | UBASH3B | 0.272 | 0.267 | 0.272 | | HMGA1 | 0.201 | 0.215 | 0.201 |
| SMYD3 | 0.398 | 0.411 | 0.398 | | NUP62CL | 0.27 | 0.495 | 0.27 | | VTI1A | 0.201 | 0.255 | 0.201 |
| GGH | 0.391 | 0.28 | 0.391 | | CCDC91 | 0.269 | 0.389 | 0.269 | | | | | |
| CENPP | 0.384 | 0.579 | 0.384 | | CFAP44 | 0.267 | 0.464 | 0.267 | | | | | |
| UXT | 0.379 | 0.26 | 0.379 | | ESCO2 | 0.266 | 0.305 | 0.266 | | | | | |
| FMN1 | 0.378 | 0.349 | 0.378 | | AMIGO2 | 0.266 | 0.37 | 0.266 | | | | | |

Supplemental Table 8. Genes upregulated in cluster F versus all other clusters at log2FC > 0.2 and p-value adjusted <= 0.05 in both cell lines

| gene | HCC1806 | MDA-MB-231 | mean_log2FC | | gene | HCC1806 | MDA-MB-231 | mean_log2FC | | gene | HCC1806 | MDA-MB-231 | mean_log2FC |
|---|---|---|---|---|---|---|---|---|---|---|---|---|---|
| L1CAM | -1.231 | -1.657 | -1.231 | | KNSTRN | -0.352 | -0.207 | -0.352 | | SLC2A6 | -0.261 | -0.484 | -0.261 |
| ATF3 | -1.222 | -1.72 | -1.222 | | MATN2 | -0.351 | -1.051 | -0.351 | | DDX6 | -0.26 | -0.274 | -0.26 |
| CREBRF | -0.902 | -0.527 | -0.902 | | ATP2B4 | -0.35 | -0.263 | -0.35 | | SLC25A25 | -0.259 | -0.211 | -0.259 |
| TCF7L1 | -0.878 | -0.733 | -0.878 | | NUP50-DT | -0.349 | -0.247 | -0.349 | | CHMP1B | -0.258 | -0.244 | -0.258 |
| TXNIP | -0.872 | -0.374 | -0.872 | | CLK4 | -0.345 | -0.265 | -0.345 | | IFIT3 | -0.258 | -2.762 | -0.258 |
| ACSF2 | -0.771 | -0.324 | -0.771 | | UBL3 | -0.343 | -0.213 | -0.343 | | CEP170B | -0.256 | -0.232 | -0.256 |
| ENSG00000259001 | -0.77 | -0.909 | -0.77 | | BHLHE40 | -0.342 | -0.724 | -0.342 | | JAK1 | -0.256 | -0.244 | -0.256 |
| YPEL5 | -0.742 | -0.27 | -0.742 | | SNAPC4 | -0.34 | -0.258 | -0.34 | | HMGCS1 | -0.256 | -0.409 | -0.256 |
| IRF1 | -0.74 | -0.798 | -0.74 | | ATG2A | -0.339 | -0.32 | -0.339 | | FASN | -0.254 | -0.336 | -0.254 |
| HEG1 | -0.737 | -0.36 | -0.737 | | ZNF117 | -0.338 | -0.508 | -0.338 | | IFIT2 | -0.254 | -3.654 | -0.254 |
| MYLIP | -0.736 | -0.201 | -0.736 | | MIIP | -0.338 | -0.591 | -0.338 | | SLC44A2 | -0.254 | -0.404 | -0.254 |
| DUSP5 | -0.73 | -0.535 | -0.73 | | BMAL1 | -0.336 | -0.488 | -0.336 | | OSBPL2 | -0.254 | -0.276 | -0.254 |
| DNAH11 | -0.71 | -1.32 | -0.71 | | TPM1 | -0.336 | -0.325 | -0.336 | | NADSYN1 | -0.254 | -0.239 | -0.254 |
| ZNF846 | -0.698 | -0.315 | -0.698 | | SLC25A29 | -0.335 | -0.316 | -0.335 | | PLEKHG3 | -0.254 | -0.271 | -0.254 |
| RMRP | -0.655 | -0.257 | -0.655 | | NFKBIA | -0.335 | -0.415 | -0.335 | | PLK1 | -0.253 | -0.303 | -0.253 |
| FBXL20 | -0.652 | -0.25 | -0.652 | | ENSG00000291065 | -0.333 | -0.281 | -0.333 | | CALM2 | -0.253 | -0.216 | -0.253 |
| MIR23AHG | -0.647 | -0.518 | -0.647 | | ADM | -0.329 | -0.426 | -0.329 | | CHASERR | -0.253 | -0.272 | -0.253 |
| DDIT4 | -0.643 | -0.324 | -0.643 | | SCNN1A | -0.328 | -0.202 | -0.328 | | NIPSNAP1 | -0.252 | -0.407 | -0.252 |
| CERCAM | -0.631 | -0.515 | -0.631 | | NUAK1 | -0.326 | -0.265 | -0.326 | | FOXP4 | -0.252 | -0.273 | -0.252 |
| TENT5A | -0.614 | -0.911 | -0.614 | | CDK19 | -0.325 | -0.216 | -0.325 | | AVPI1 | -0.25 | -0.395 | -0.25 |
| FN1 | -0.589 | -0.222 | -0.589 | | NFKB2 | -0.324 | -0.468 | -0.324 | | HMGCR | -0.248 | -0.292 | -0.248 |
| PLA2R1 | -0.584 | -0.426 | -0.584 | | ZFP36L2 | -0.324 | -0.888 | -0.324 | | PANX1 | -0.248 | -0.642 | -0.248 |
| RAB11FIP1 | -0.569 | -0.499 | -0.569 | | EEIG1 | -0.323 | -0.444 | -0.323 | | SLC29A2 | -0.247 | -0.26 | -0.247 |
| KDM5B | -0.561 | -0.388 | -0.561 | | NEK2 | -0.323 | -0.408 | -0.323 | | PITPNM3 | -0.247 | -0.32 | -0.247 |
| NEDD9 | -0.547 | -1.293 | -0.547 | | CRY2 | -0.322 | -0.413 | -0.322 | | ZNF558 | -0.246 | -0.25 | -0.246 |
| TUFT1 | -0.538 | -0.573 | -0.538 | | PAPSS2 | -0.322 | -0.226 | -0.322 | | ZSCAN29 | -0.246 | -0.296 | -0.246 |
| HBP1 | -0.526 | -0.387 | -0.526 | | ZFAS1 | -0.322 | -0.234 | -0.322 | | PIK3C3 | -0.245 | -0.219 | -0.245 |
| OSMR-DT | -0.522 | -0.313 | -0.522 | | PRTG | -0.32 | -0.352 | -0.32 | | RNF5 | -0.244 | -0.207 | -0.244 |
| RBKS | -0.507 | -0.303 | -0.507 | | EAF1 | -0.32 | -0.218 | -0.32 | | SREBF2 | -0.243 | -0.27 | -0.243 |
| MKNK2 | -0.506 | -0.297 | -0.506 | | TRIM59 | -0.32 | -0.243 | -0.32 | | PCYT2 | -0.243 | -0.38 | -0.243 |
| FSTL1 | -0.498 | -0.258 | -0.498 | | SPSB1 | -0.319 | -0.242 | -0.319 | | NCK2 | -0.241 | -0.239 | -0.241 |
| DUSP16 | -0.497 | -0.306 | -0.497 | | MAFK | -0.317 | -0.447 | -0.317 | | RHBDF2 | -0.241 | -0.368 | -0.241 |
| OPTN | -0.492 | -0.49 | -0.492 | | NDRG1 | -0.317 | -0.276 | -0.317 | | EPS8L2 | -0.24 | -0.406 | -0.24 |
| LINC00370 | -0.49 | -0.358 | -0.49 | | LEPR | -0.316 | -0.287 | -0.316 | | CCSAP | -0.239 | -0.448 | -0.239 |
| DBN1 | -0.481 | -0.247 | -0.481 | | BTN3A3 | -0.314 | -0.733 | -0.314 | | RHOQ | -0.238 | -0.372 | -0.238 |
| DNAJC6 | -0.478 | -0.374 | -0.478 | | IL18 | -0.314 | -0.314 | -0.314 | | TAPBP | -0.237 | -0.322 | -0.237 |
| PRKCD | -0.475 | -0.233 | -0.475 | | POMT1 | -0.313 | -0.25 | -0.313 | | TAF8 | -0.232 | -0.204 | -0.232 |
| HIVEP2 | -0.473 | -0.527 | -0.473 | | FBXL12 | -0.312 | -0.22 | -0.312 | | SIRT7 | -0.231 | -0.277 | -0.231 |
| C6orf132 | -0.463 | -0.879 | -0.463 | | PIMREG | -0.312 | -0.257 | -0.312 | | SLC45A4 | -0.23 | -0.404 | -0.23 |
| BCAS4 | -0.461 | -0.377 | -0.461 | | ARRDC3 | -0.311 | -0.422 | -0.311 | | SORBS3 | -0.229 | -0.35 | -0.229 |
| SCD | -0.46 | -0.55 | -0.46 | | ARHGAP45 | -0.31 | -0.873 | -0.31 | | BCL2L11 | -0.228 | -0.263 | -0.228 |
| ARL6IP1 | -0.458 | -0.397 | -0.458 | | IFNGR2 | -0.309 | -0.327 | -0.309 | | CDV3 | -0.228 | -0.272 | -0.228 |
| RNF19B | -0.457 | -0.38 | -0.457 | | B3GNT2 | -0.308 | -0.285 | -0.308 | | DSP | -0.228 | -0.414 | -0.228 |
| RAB11FIP5 | -0.457 | -0.388 | -0.457 | | KIF20A | -0.308 | -0.463 | -0.308 | | PACSIN3 | -0.228 | -0.281 | -0.228 |
| IL10RB | -0.44 | -0.266 | -0.44 | | ENSG00000229618 | -0.307 | -0.63 | -0.307 | | B4GALT5 | -0.228 | -0.633 | -0.228 |
| SNX33 | -0.437 | -0.306 | -0.437 | | ZNF724.1 | -0.304 | -0.296 | -0.304 | | DEPDC1 | -0.228 | -0.239 | -0.228 |
| HINFP | -0.436 | -0.221 | -0.436 | | LATS2 | -0.303 | -0.381 | -0.303 | | TMEM50B | -0.228 | -0.262 | -0.228 |
| UBE2Q2P1 | -0.434 | -0.303 | -0.434 | | MAP3K11 | -0.302 | -0.3 | -0.302 | | RPL12 | -0.227 | -0.228 | -0.227 |
| TRIM65 | -0.432 | -0.421 | -0.432 | | PDXDC2P | -0.298 | -0.277 | -0.298 | | VPS11 | -0.227 | -0.221 | -0.227 |
| IFIT1 | -0.432 | -2.904 | -0.432 | | HIF1A-AS3 | -0.296 | -0.3 | -0.296 | | BAHCC1 | -0.227 | -0.384 | -0.227 |
| IQSEC1 | -0.431 | -0.241 | -0.431 | | COL6A1 | -0.295 | -0.239 | -0.295 | | ATP8B1 | -0.227 | -0.339 | -0.227 |
| ZNFX1 | -0.428 | -0.658 | -0.428 | | ZFYVE27 | -0.295 | -0.209 | -0.295 | | RAB11FIP3 | -0.226 | -0.332 | -0.226 |
| PLAC8 | -0.427 | -0.312 | -0.427 | | UBXN7 | -0.291 | -0.239 | -0.291 | | ANKZF1 | -0.225 | -0.402 | -0.225 |
| PECR | -0.426 | -0.357 | -0.426 | | TNKS1BP1 | -0.29 | -0.259 | -0.29 | | FAM234A | -0.225 | -0.283 | -0.225 |
| SLC22A5 | -0.425 | -0.601 | -0.425 | | ARHGAP27 | -0.29 | -0.278 | -0.29 | | TBC1D10A | -0.224 | -0.523 | -0.224 |
| BCL6 | -0.424 | -0.288 | -0.424 | | SNHG12 | -0.288 | -0.25 | -0.288 | | EIF2AK3 | -0.223 | -0.343 | -0.223 |
| MINDY2 | -0.423 | -0.282 | -0.423 | | GRN | -0.287 | -0.218 | -0.287 | | TNFAIP1 | -0.223 | -0.349 | -0.223 |
| CPEB4 | -0.422 | -0.241 | -0.422 | | RIT1 | -0.286 | -0.315 | -0.286 | | MIR3936HG | -0.222 | -0.385 | -0.222 |
| UCP2 | -0.422 | -0.332 | -0.422 | | MELTF | -0.284 | -0.28 | -0.284 | | CHST3 | -0.222 | -0.222 | -0.222 |
| RIPK4 | -0.421 | -0.505 | -0.421 | | IFNAR2 | -0.283 | -0.294 | -0.283 | | MAP7D1 | -0.22 | -0.28 | -0.22 |
| SREBF1 | -0.419 | -0.396 | -0.419 | | CCN1 | -0.282 | -0.259 | -0.282 | | CPEB3 | -0.22 | -0.215 | -0.22 |
| PPARA | -0.415 | -0.215 | -0.415 | | RIPOR1 | -0.281 | -0.295 | -0.281 | | VPS26C | -0.219 | -0.202 | -0.219 |
| ETV3 | -0.413 | -0.344 | -0.413 | | LRRC37B | -0.281 | -0.247 | -0.281 | | PLD2 | -0.219 | -0.335 | -0.219 |
| MGAT4A | -0.413 | -0.435 | -0.413 | | MTUS1 | -0.28 | -0.478 | -0.28 | | WASHC2C | -0.219 | -0.225 | -0.219 |
| MTCL2 | -0.411 | -0.21 | -0.411 | | PSMG4 | -0.279 | -0.42 | -0.279 | | NAT9 | -0.213 | -0.291 | -0.213 |
| TNFAIP2 | -0.403 | -0.398 | -0.403 | | DRAM1 | -0.279 | -0.389 | -0.279 | | C1orf52 | -0.213 | -0.243 | -0.213 |
| CLCN7 | -0.401 | -0.285 | -0.401 | | DNASE2 | -0.279 | -0.329 | -0.279 | | CD55 | -0.213 | -0.422 | -0.213 |
| HIP1R | -0.399 | -0.572 | -0.399 | | RSRP1 | -0.278 | -0.344 | -0.278 | | PRKCZ | -0.212 | -0.939 | -0.212 |
| MXI1 | -0.397 | -0.506 | -0.397 | | PARD6B | -0.277 | -0.468 | -0.277 | | NTN4 | -0.211 | -0.314 | -0.211 |
| PLEKHM2 | -0.396 | -0.21 | -0.396 | | RHOF | -0.276 | -0.269 | -0.276 | | MXRA7 | -0.21 | -0.398 | -0.21 |
| CENPF | -0.396 | -0.237 | -0.396 | | CCNB1 | -0.276 | -0.477 | -0.276 | | RP9P | -0.209 | -0.533 | -0.209 |
| MAFF | -0.393 | -0.339 | -0.393 | | BAG1 | -0.274 | -0.262 | -0.274 | | KLHL36 | -0.209 | -0.312 | -0.209 |
| CKAP2 | -0.387 | -0.283 | -0.387 | | NSMF | -0.274 | -0.459 | -0.274 | | CASZ1 | -0.208 | -0.312 | -0.208 |
| HERPUD1 | -0.383 | -0.231 | -0.383 | | SDC4 | -0.273 | -0.315 | -0.273 | | MGAT4B | -0.207 | -0.334 | -0.207 |
| ITGA5 | -0.383 | -0.228 | -0.383 | | SAMD9L | -0.273 | -1.171 | -0.273 | | LLGL2 | -0.207 | -0.908 | -0.207 |
| SLC1A3 | -0.381 | -0.916 | -0.381 | | CDK9 | -0.27 | -0.292 | -0.27 | | HMGB2 | -0.207 | -0.253 | -0.207 |
| ARID4A | -0.379 | -0.533 | -0.379 | | ARHGAP23 | -0.27 | -0.317 | -0.27 | | F11R | -0.205 | -0.415 | -0.205 |
| RNF207 | -0.376 | -0.314 | -0.376 | | SMOX | -0.269 | -0.553 | -0.269 | | ECHDC2 | -0.205 | -0.225 | -0.205 |
| ELOVL7 | -0.373 | -0.656 | -0.373 | | MUS81 | -0.268 | -0.234 | -0.268 | | PER3 | -0.205 | -0.217 | -0.205 |
| TESK2 | -0.369 | -0.35 | -0.369 | | UBTD1 | -0.266 | -0.305 | -0.266 | | LSS | -0.204 | -0.365 | -0.204 |
| HELZ2 | -0.367 | -0.337 | -0.367 | | BLVRA | -0.265 | -0.322 | -0.265 | | CCDC9 | -0.202 | -0.412 | -0.202 |
| TOB1 | -0.367 | -0.382 | -0.367 | | CCDC186 | -0.265 | -0.368 | -0.265 | | ANXA4 | -0.202 | -0.282 | -0.202 |

| | | | | | | | | | | | | | |
|---|---|---|---|---|---|---|---|---|---|---|---|---|---|
| APOL6 | -0.366 | -1.101 | -0.366 | | ANKRD16 | -0.264 | -0.309 | -0.264 | | BAG3 | -0.2 | -0.218 | -0.2 |
| CCNG2 | -0.364 | -0.526 | -0.364 | | KLC4 | -0.263 | -0.306 | -0.263 | | ENSG00000 293331 | -0.2 | -0.213 | -0.2 |
| FBXO17 | -0.362 | -0.25 | -0.362 | | AGR2 | -0.263 | -0.808 | -0.263 | | | | | |
| KCNQ1OT1 | -0.361 | -0.352 | -0.361 | | EPAS1 | -0.262 | -0.565 | -0.262 | | | | | |
| TDRD7 | -0.359 | -0.517 | -0.359 | | BTN3A1 | -0.262 | -0.574 | -0.262 | | | | | |
| AURKA | -0.358 | -0.413 | -0.358 | | ENSG00000 250764 | -0.262 | -0.376 | -0.262 | | | | | |

Supplemental Table 9. Genes downregulated in cluster F versus all other clusters at log2FC < -0.2 and p-value adjusted <= 0.05 in both cell lines

| gene | HCC1806 | MDA-MB-231 | mean_log2FC | | gene | HCC1806 | MDA-MB-231 | mean_log2FC |
|---|---|---|---|---|---|---|---|---|
| MT2A | 0.761 | 0.622 | 0.761 | | MICOS13 | 0.293 | 0.394 | 0.293 |
| ADIRF | 0.716 | 0.818 | 0.716 | | SNRPG | 0.29 | 0.275 | 0.29 |
| NAALADL2 | 0.715 | 0.56 | 0.715 | | LPAR1 | 0.29 | 0.584 | 0.29 |
| CDKN3 | 0.636 | 0.289 | 0.636 | | RMI1 | 0.287 | 0.206 | 0.287 |
| COL13A1 | 0.6 | 0.517 | 0.6 | | ZNF155 | 0.286 | 0.316 | 0.286 |
| ADAMTS6 | 0.564 | 0.898 | 0.564 | | RPS26 | 0.28 | 0.203 | 0.28 |
| DCTN3 | 0.549 | 0.269 | 0.549 | | ARPC2 | 0.279 | 0.229 | 0.279 |
| RPS21 | 0.549 | 0.255 | 0.549 | | CKAP2 | 0.279 | 0.225 | 0.279 |
| KIZ-AS1 | 0.547 | 0.386 | 0.547 | | ENSG00000271727 | 0.277 | 0.466 | 0.277 |
| RPS27L | 0.539 | 0.225 | 0.539 | | IFI27L1 | 0.276 | 0.242 | 0.276 |
| MRPS33 | 0.522 | 0.253 | 0.522 | | SAMD1 | 0.27 | 0.373 | 0.27 |
| ZNF302 | 0.498 | 0.323 | 0.498 | | TMBIM4 | 0.268 | 0.286 | 0.268 |
| HCFC1R1 | 0.497 | 0.547 | 0.497 | | FRMD5 | 0.268 | 0.291 | 0.268 |
| HMGN5 | 0.492 | 1.236 | 0.492 | | NFU1 | 0.263 | 0.281 | 0.263 |
| NDUFC1 | 0.488 | 0.354 | 0.488 | | YEATS4 | 0.262 | 0.233 | 0.262 |
| IRF2BPL | 0.479 | 0.386 | 0.479 | | ARHGEF39 | 0.261 | 0.358 | 0.261 |
| SPTLC3 | 0.458 | 0.376 | 0.458 | | EXTL3 | 0.257 | 0.359 | 0.257 |
| TMEM106C | 0.455 | 0.249 | 0.455 | | DNAJC19 | 0.256 | 0.335 | 0.256 |
| CHMP2A | 0.447 | 0.34 | 0.447 | | ACP1 | 0.245 | 0.274 | 0.245 |
| RPSA2 | 0.443 | 0.278 | 0.443 | | PCBP1 | 0.241 | 0.217 | 0.241 |
| H4C3 | 0.442 | 0.371 | 0.442 | | GAS2L3 | 0.234 | 0.276 | 0.234 |
| CUTA | 0.442 | 0.42 | 0.442 | | TMEM14B | 0.233 | 0.348 | 0.233 |
| CKLF | 0.442 | 0.307 | 0.442 | | EHF | 0.225 | 0.578 | 0.225 |
| RPS20 | 0.416 | 0.283 | 0.416 | | UPF3B | 0.222 | 0.359 | 0.222 |
| NR2F2-AS1 | 0.416 | 0.561 | 0.416 | | GALM | 0.221 | 0.482 | 0.221 |
| NDUFB1 | 0.416 | 0.358 | 0.416 | | SERTAD2 | 0.22 | 0.339 | 0.22 |
| RIBC2 | 0.415 | 0.381 | 0.415 | | MBOAT1 | 0.219 | 0.604 | 0.219 |
| NRM | 0.402 | 0.399 | 0.402 | | ETAA1 | 0.219 | 0.32 | 0.219 |
| ZNF395 | 0.389 | 0.308 | 0.389 | | CFAP36 | 0.218 | 0.235 | 0.218 |
| ARHGDIB | 0.382 | 0.353 | 0.382 | | TPGS2 | 0.215 | 0.323 | 0.215 |
| MTIF3 | 0.376 | 0.248 | 0.376 | | ARHGAP18 | 0.213 | 0.428 | 0.213 |
| NDUFB4 | 0.372 | 0.26 | 0.372 | | FAM83D | 0.21 | 0.233 | 0.21 |
| ENSG00000255094 | 0.369 | 0.618 | 0.369 | | NUBP1 | 0.209 | 0.316 | 0.209 |
| FTL | 0.353 | 0.479 | 0.353 | | FOXM1 | 0.208 | 0.208 | 0.208 |
| ELP4 | 0.32 | 0.219 | 0.32 | | TUBA1B | 0.208 | 0.306 | 0.208 |
| NR2F2 | 0.315 | 0.608 | 0.315 | | NQO1 | 0.204 | 0.452 | 0.204 |
| PCGF3-AS1 | 0.31 | 0.232 | 0.31 | | DPH6 | 0.203 | 0.429 | 0.203 |
| CREM | 0.309 | 0.27 | 0.309 | | PNRC2 | 0.203 | 0.315 | 0.203 |
| ASB8 | 0.302 | 0.419 | 0.302 | | CRIPT | 0.201 | 0.284 | 0.201 |

Supplemental Table 10. Genes upregulated in fusion vs. control cells at log2FC > 0.2 and p-value adjusted <= 0.05 in both cell lines

| gene | HCC1806 | MDA-MB-231 | mean_log2FC | | gene | HCC1806 | MDA-MB-231 | mean_log2FC |
|---|---|---|---|---|---|---|---|---|
| CADPS2 | -1.397 | -0.581 | -1.397 | | PEX19 | -0.277 | -0.279 | -0.277 |
| PDE3A | -1.254 | -1.106 | -1.254 | | TRPV1 | -0.277 | -0.265 | -0.277 |
| PRR5L | -0.856 | -0.848 | -0.856 | | HS6ST1 | -0.276 | -0.575 | -0.276 |
| TBC1D4 | -0.552 | -0.361 | -0.552 | | CHKA | -0.276 | -0.477 | -0.276 |
| PPP1R16A | -0.467 | -0.448 | -0.467 | | ENSG00000261786 | -0.275 | -0.362 | -0.275 |
| AASS | -0.421 | -0.466 | -0.421 | | DDAH1 | -0.267 | -0.523 | -0.267 |
| BAHCC1 | -0.399 | -0.508 | -0.399 | | UPP1 | -0.267 | -0.373 | -0.267 |
| MAFK | -0.382 | -0.393 | -0.382 | | NUAK1 | -0.266 | -0.695 | -0.266 |
| ZNF707 | -0.372 | -0.238 | -0.372 | | SLC22A5 | -0.264 | -0.578 | -0.264 |
| HEG1 | -0.372 | -0.596 | -0.372 | | SLC44A2 | -0.254 | -0.49 | -0.254 |
| MT-RNR1 | -0.362 | -0.22 | -0.362 | | LLGL2 | -0.247 | -1.519 | -0.247 |
| LRP5L | -0.354 | -0.528 | -0.354 | | TUFT1 | -0.243 | -0.661 | -0.243 |
| SLC25A33 | -0.344 | -0.296 | -0.344 | | FASN | -0.237 | -0.631 | -0.237 |
| DBN1 | -0.331 | -0.453 | -0.331 | | ZPR1 | -0.235 | -0.345 | -0.235 |
| TSEN54 | -0.327 | -0.394 | -0.327 | | BYSL | -0.231 | -0.326 | -0.231 |
| RRP9 | -0.317 | -0.384 | -0.317 | | SORT1 | -0.223 | -0.267 | -0.223 |
| ARHGAP27 | -0.301 | -0.541 | -0.301 | | NT5DC3 | -0.215 | -0.37 | -0.215 |
| PFKFB2 | -0.3 | -0.436 | -0.3 | | SPON1-AS1 | -0.209 | -0.435 | -0.209 |
| FBRS | -0.298 | -0.26 | -0.298 | | MTCL1 | -0.204 | -0.341 | -0.204 |
| F2RL1 | -0.292 | -0.33 | -0.292 | | | | | |

Supplemental Table 11. Genes downregulated in fusion vs. control cells at log2FC < -0.2 and p-value adjusted <= 0.05 in both cell lines
